# Multi-omics profiling of U2AF1 mutants dissects pathogenic mechanisms affecting RNA granules in myeloid malignancies

**DOI:** 10.1101/2021.04.22.441020

**Authors:** Giulia Biancon, Poorval Joshi, Joshua T Zimmer, Torben Hunck, Yimeng Gao, Mark D Lessard, Edward Courchaine, Andrew ES Barentine, Martin Machyna, Valentina Botti, Ashley Qin, Rana Gbyli, Amisha Patel, Yuanbin Song, Lea Kiefer, Gabriella Viero, Nils Neuenkirchen, Haifan Lin, Joerg Bewersdorf, Matthew D Simon, Karla M Neugebauer, Toma Tebaldi, Stephanie Halene

**Affiliations:** Section of Hematology, Department of Internal Medicine and Yale Comprehensive Cancer Center, Yale University School of Medicine, New Haven, CT, USA; Department of Molecular Biophysics and Biochemistry, Yale University School of Medicine, New Haven, CT, USA; Department of Cell Biology, Yale University School of Medicine, New Haven, CT, USA; Department of Biomedical Engineering, Yale University, New Haven, CT, USA; Department of Hematologic Oncology, Sun Yat-sen University Cancer Center, State Key Laboratory of Oncology in South China, Collaborative Innovation Center for Cancer Medicine, Guangzhou, China; University of California San Francisco, San Francisco, CA, USA; Institute of Biophysics, CNR, Trento, Italy; Yale Stem Cell Center, Yale University School of Medicine, New Haven, CT, USA; Department of Cellular, Computational and Integrative Biology (CIBIO), University of Trento, Trento, Italy

## Abstract

Somatic mutations in splicing factors are of significant interest in myeloid malignancies and other cancers. U2AF1, together with U2AF2, is essential for 3’ splice site recognition. U2AF1 mutations result in aberrant splicing, but the molecular mechanism and the full spectrum of consequences on RNA biology have not been fully elucidated to date. We performed multi-omics profiling of *in vivo* RNA binding, splicing and turnover for U2AF1 S34F and Q157R mutants. We dissected specific binding signals of U2AF1 and U2AF2 and showed that U2AF1 mutations individually alter U2AF1-RNA binding, resulting in defective U2AF2 recruitment. We demonstrated a complex relationship between differential binding and splicing, expanding upon the currently accepted loss-of-binding model. Finally, we observed that U2AF1 mutations increase the formation of stress granules in both cell lines and primary acute myeloid leukemia samples. Our results uncover U2AF1 mutation-dependent pathogenic RNA mechanisms and provide the basis for developing targeted therapeutic strategies.

## Introduction

Somatic mutations in splicing factor genes SF3B1, SRSF2, U2AF1 and ZRSR2 function as drivers of myeloid malignancies and solid cancers such as lung adenocarcinoma^1,2^. They are generally mutually exclusive and associated with clinical outcome^3,4^. They occur in approximately 10% of *de novo* acute myeloid leukemia (AML) while reaching >50% in myelodysplastic syndromes (MDS) and secondary AML^5–7^. MDS and AML are clonal hematopoietic stem cell malignancies characterized by bone marrow failure and peripheral blood cytopenias due to the uncontrolled proliferation and dysfunctional maturation of myeloid blasts^8,9^. Despite novel agents leading to significant improvements in patient outcome^10,11^, the 5-year relative survival rate is only 38.3% in MDS and 28.7% in AML^12^, highlighting the need to better understand pathogenic mechanisms for the development of more efficient targeted therapies.

U2AF1 (U2 small nuclear RNA auxiliary factor 1), together with U2AF2, participates in pre-mRNA splicing by forming the U2AF complex that recognizes the 3’ splice site (3’SS) of U2 introns (**Figure 1A**) and recruits U2 small nuclear ribonucleoproteins (snRNPs)^13–16^. U2AF1 domains include a U2AF homology motif (UHM) that mediates the heterodimerization with U2AF2^17^, and two conserved CCCH-type zinc fingers (ZnF1 and ZnF2) that cooperatively bind the 3’SS intron-exon boundaries of target RNA^18^. Heterozygous hotspot mutations affecting the S34 and Q157 residues within the ZnF1 and ZnF2 domains, respectively, result in sequence-dependent mis-splicing of genes crucial for hematopoiesis^19,20^. It has been previously reported that the S34 mutation preferentially leads to exclusion of exons bearing a U in position −3 of the 3’SS region and to inclusion of exons bearing a C at the same position. The Q157 mutation preferentially excludes exons starting with A and includes exons starting with G in position +1 of the 3’SS region ^21,22^. Yet the molecular mechanisms leading to these splicing alterations and ultimately to disease are still not fully understood.

**Figure 1.**
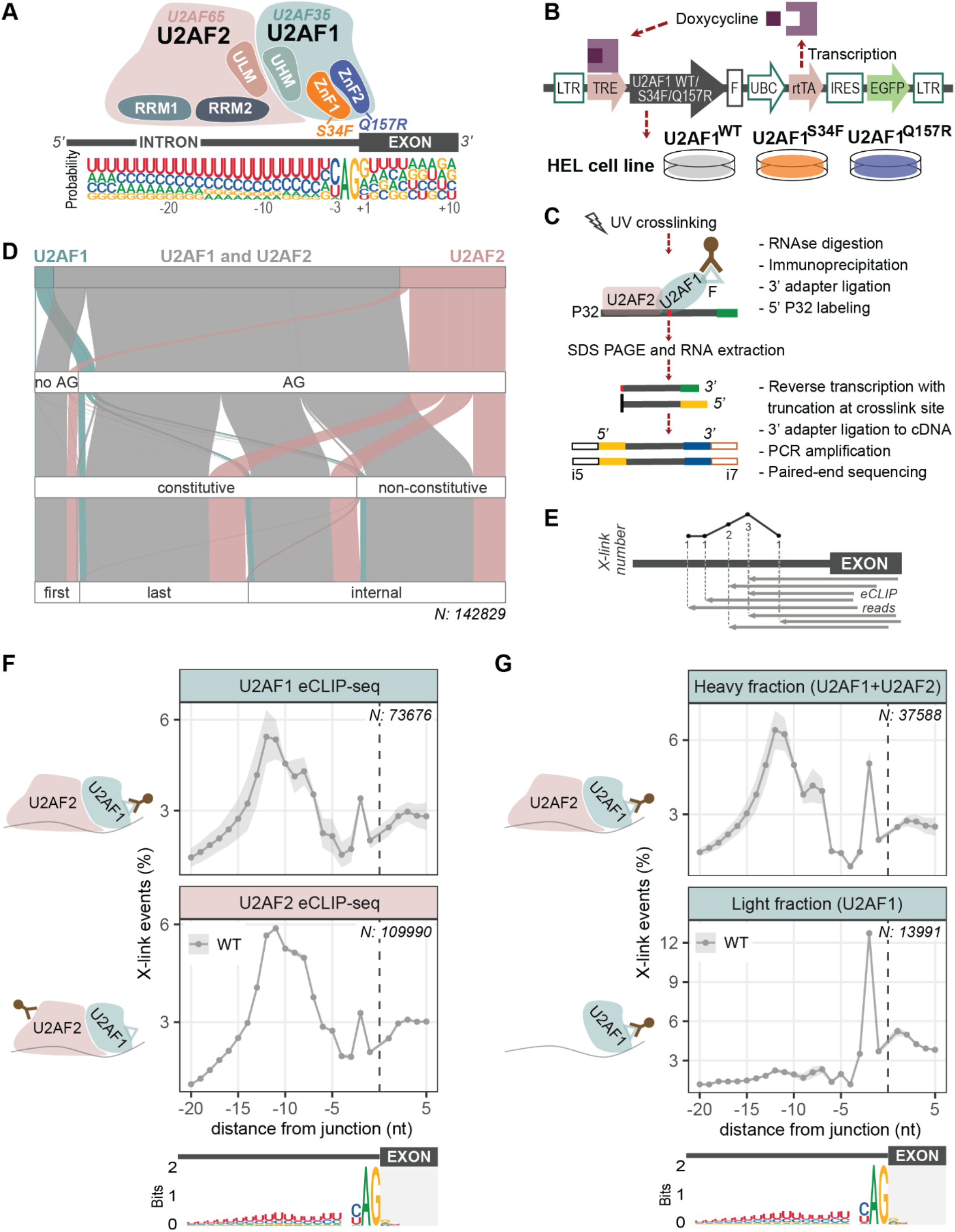
In-depth mapping of U2AF-RNA binding reveals the in vivo position of the U2AF1 subunit within the 3’SS. **(A)** Schematic representation of the U2AF complex on the intronic 3’SS. The presumed locations of U2AF1 and U2AF2, their relevant domains, the approximate position of U2AF1 pathological mutations and the consensus sequence of the human 3’SS are displayed. RRM, RNA recognition motif; ULM, U2AF ligand motif; UHM, U2AF homology motif; ZnF, zinc finger domain. **(B)** Human erythroleukemia (HEL) cells expressing FLAG-tagged WT, S34F or Q157R U2AF1 after doxycycline induction. **(C)** Schematic overview of the U2AF1 eCLIP-seq protocol. **(D)** Alluvial plot showing different classes of intron-exon junctions bound by U2AF1, U2AF2 or both (color-coded), based on eCLIP-seq. Junction classes are defined by the presence of canonical AG at the intronic 3’SS, constitutive *vs* non-constitutive exon, and the position within the transcript. **(E)** Illustrative intron-exon junction where eCLIP-seq coverage is quantified with single-nucleotide resolution based on the number of crosslinking events. **(F)** Binding metaprofile (mean±SEM of the percentage of crosslinking events) and 3’SS sequence logo for U2AF1 (top panel, n=3) and U2AF2 (bottom panel, n=4) eCLIP-seq. N, number of intron-exon junctions. **(G)** Binding metaprofile (mean±SEM) and 3’SS sequence logo for heavy (top panel, n=4) and light (bottom panel, n=3) fractions of U2AF1 WT freCLIP-seq. N, number of intron-exon junctions.

Previous studies, relying on *in vitro* binding assays and structural predictions, suggest that aberrant splicing may directly result from loss-of-binding, where U2AF1 mutants display lower affinity for splice junctions of excluded exons or retained introns^21,23,24^. However, recent structural studies by Yoshida *et al.*^25^ show that the S34F/Y mutants do not display reduced binding affinities for 3’SS sequences that predict exon exclusion, suggesting the presence of a more complex and not yet dissected binding-splicing relationship.

We here present multi-omics integration of the RNA interactome, transcriptome and RNA turnover in human erythroleukemia (HEL) cells expressing wild-type (WT) or mutant (S34F and Q157R) U2AF1.

Enhanced UV-crosslinking and immunoprecipitation followed by sequencing (eCLIP-seq)^26^ allows the determination of protein-RNA binding at single-nucleotide resolution^27^ with improvements in sensitivity and sequencing depth in comparison to previous CLIP-seq methods^28,29^. By developing a fractionated (fr)eCLIP-seq protocol, we were able to dissect U2AF1 from U2AF2 RNA binding signals within the 3’SS region *in vivo*. Moreover, we identified specific changes in mutant U2AF1 binding. Specifically, we observed a *de novo* binding peak in position −3 for S34F and in position +1 for Q157R, matching exactly the critical positions identified by differential splicing analysis on RNA-seq data. Surprisingly, combined binding-splicing analysis showed that, while the Q157R mutant predominantly exhibited loss-of-binding, the S34F mutant exhibited a gain-of-binding pattern, where splicing progression was impaired upon increased mutant binding.

Finally, we report that both U2AF1 mutants altered RNA granule biology, specifically affecting the abundance and composition of stress granule (SG)-enriched transcripts and proteins. RNA turnover analysis using TimeLapse (TL)-seq and immunofluorescence (IF) imaging confirmed enhanced SG formation in U2AF1-mutant cell lines and primary AML cells.

Collectively, our results provide in-depth understanding of the *in vivo* pathogenic molecular mechanisms conferred by U2AF1 mutations in myeloid malignancies, uncovering novel potential therapeutic vulnerabilities.

## Results

### Fractionated eCLIP-seq isolates individual binding of U2AF1 from U2AF2 in vivo

To investigate the role of U2AF1 mutants (**Figure 1A**) in myeloid malignancies, we generated HEL cell lines with doxycycline (dox)-inducible expression of FLAG-tagged WT, S34F or Q157R U2AF1 via lentiviral transduction^5^ (**Figure 1B**). We sorted GFP^+^ cells and induced FLAG-tagged U2AF1 proteins for 48 hours. Inducible expression was verified by RT-PCR followed by Sanger sequencing and quantitated by western blot. After doxycycline induction, HEL cells stably expressed FLAG-tagged U2AF1 to levels only modestly above baseline (**Figures S1A-S1B**).

To characterize transcriptome-wide *in vivo* U2AF-RNA interactions, we first performed U2AF1 and U2AF2 eCLIP-seq in U2AF1 WT cells (**Table S1**). Crosslinked exogenous U2AF1-RNA complexes were isolated by immunoprecipitation with an anti-FLAG antibody^30^ (**Figure 1C**), while endogenous U2AF2-RNA complexes were isolated with an anti-U2AF2 antibody^31^.

Selecting a window ranging from −30 to +5 nucleotides around the 3’SS, we identified 142,829 intron-exon junctions in 10,821 genes specifically bound by U2AF1 (n=5,670), U2AF2 (n=32,097) or both (n=105,062) (**Figure 1D**). With approximately 73.6% of the identified junctions bound by the heterodimer, our deep characterization demonstrated the cooperative function of U2AF1 and U2AF2 in ensuring inclusion of constitutive exons and regulation of alternative splicing of non-constitutive exons. The U2AF complex recognized internal and last exons with similar efficiency (54.1% and 36.2%, respectively), but it was also involved in the binding of first exons, confirming possible non-canonical roles of U2AF1 in translation regulation^32^. Consistent with the published literature^2,33^, U2AF1 and U2AF2 mainly bound AG-dependent 3’SS (90.8% of the identified junctions) (**Figure 1D**).

To dissect RNA binding at single-nucleotide resolution, we developed a computational binding analysis pipeline based on the interpretation of eCLIP-seq data reported by Van Nostrand *et al.*^26^. Since protein-RNA crosslinks stop reverse transcription, the nucleotide right after the end of the sequenced read corresponds to the originally crosslinked nucleotide; this allows to map and quantify interactions at each nucleotide (**Figure 1E**).

Considering internal and last exons of protein coding genes, we built 3’SS binding metaprofiles for 73,676 intron-exon junctions in U2AF1 eCLIP-seq and 109,990 junctions in U2AF2 eCLIP-seq (**Figure 1F**). These metaprofiles showed a broad peak over the −15;-5 region, corresponding to the polypyrimidine tract (PPT), and a sharp peak at the −2 nucleotide. The near-identical shapes for U2AF1 and U2AF2 binding metaprofiles demonstrated that U2AF1 and U2AF2 mostly bind their target RNAs as a dimer and standard eCLIP-seq could not resolve their individual binding contributions.

To dissect U2AF1 *vs* U2AF2 specific binding, we developed fractionated eCLIP-seq by size-selection of membrane regions to isolate the “light fraction”, containing the U2AF1 monomer plus bound RNA, and the “heavy fraction”, including the U2AF1-U2AF2 heterodimer plus bound RNA (**Figure S1C**, **Table S1**). Analysis of single junctions and generation of metaprofiles of the light and heavy fractions allowed to isolate *in vivo* binding of U2AF1 from the U2AF complex. The U2AF1 peak was precisely located in position −2 of the 3’SS, corresponding to the A of the conserved AG dinucleotide, while the peak over the PPT due to U2AF2 binding was absent in the light fraction (**Figure 1G**, **Figure S1D**).

Therefore, we mapped U2AF interactions across ~150,000 splice junctions in 11,000 human genes and we were able to specifically localize and dissect for the first time *in vivo* binding of U2AF1 and U2AF2.

### S34F and Q157R mutations alter U2AF1-RNA binding in specific positions relevant to differential splicing

To understand whether U2AF1 mutations alter RNA binding we applied freCLIP-seq to cells expressing U2AF1 S34F or Q157R (**Figure S1C**). We confirmed successful separation of the binding signals of U2AF1 monomer (light fraction, main peak in position −2) and U2AF heterodimer (heavy fraction) (**Figure 2A**). Importantly, metaprofiles pointed to distinct conformational changes in the binding of both U2AF1 mutants compared to WT U2AF1. We observed *de novo* peaks in position −3 of the 3’SS region for U2AF1 S34F and in position +1 for U2AF1 Q157R (**Figure 2A**). Interestingly, analysis of the heavy fraction metaprofiles revealed a relative reduction of the signal over the polypyrimidine track, suggesting that U2AF1 mutations altered U2AF2-RNA binding (**Figure 2A**, upper panel).

**Figure 2.**
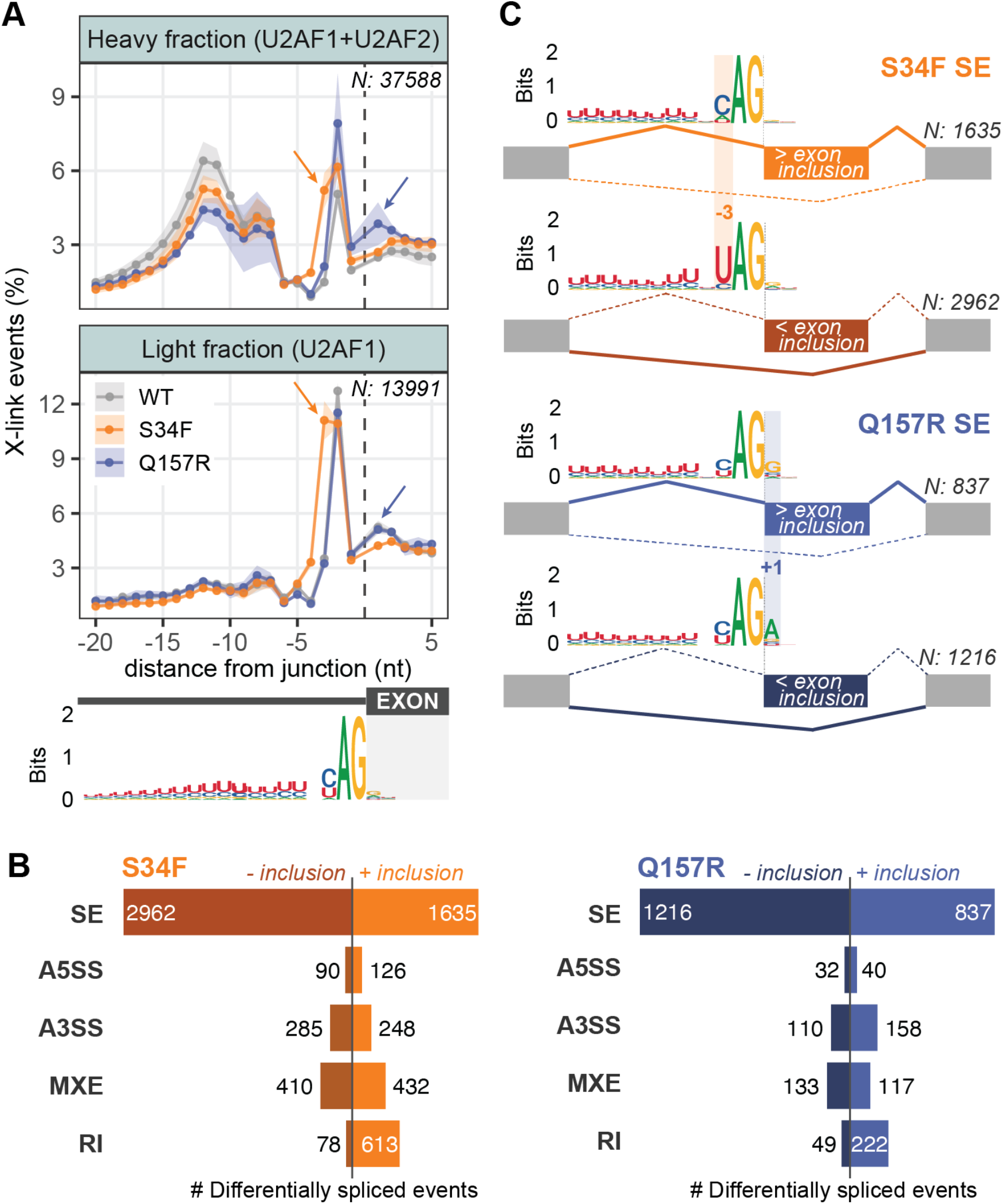
S34F and Q157R mutations cause U2AF1 binding changes matching differentially spliced sequence-specific positions. **(A)** Binding metaprofiles (mean±SEM of the percentage of crosslinking events) and 3’SS sequence logo for heavy (top panel) and light (bottom panel) fractions of U2AF1 freCLIP-seq, comparing WT with S34F (n=3) and Q157R (n=2) mutations. Arrows indicate the emergent binding peaks in position −3 for S34F mutant and +1 for Q157R mutant, respectively. N, number of intron-exon junctions. **(B)** Number of differentially spliced events in S34F and Q157R compared with WT (absolute delta PSI>10%; FDR<0.05). SE, skipped exons; A5SS, alternative 5’SS; A3SS, alternative 3’SS, MXE, mutually exclusive exons; RI, retained introns. **(C)** 3’SS sequence logos for differential SE events in U2AF1 S34F (top panel) and Q157R (bottom panel) conditions. Sequence-specific positions between more included and less included exons are highlighted (position −3 for S34F mutant, +1 for Q157R mutant) N, number of differentially spliced events.

To explore the relationship between binding and splicing perturbations caused by U2AF1 mutations, we generated RNA-seq libraries from dox-induced and uninduced U2AF1 WT, S34F and Q157R cells. After exclusion of solely dox-dependent events (see Methods; **Figure S2A**), alternative splicing analysis of S34F and Q157R mutant *vs* WT cells detected 6,879 and 2,914 differentially spliced events, respectively, with a difference in Percent Spliced-In (delta PSI) > 10% and False Discovery Rate (FDR) < 0.05 (**Figure 2B; Table S2**). Skipped exons (SE) represented the most frequent event type, with a trend favoring exon exclusion over exon inclusion in both mutants. We also observed differences in alternative 3’SS (A3SS; 7.7% of total events in S34F, 9.2% in Q157R), mutually exclusive exons (MXE; 12.2% in S34F, 8.6% in Q157R) and retained introns (RI; 10.0% in S34F, 9.3% in Q157R), with a trend favoring intron retention in both mutants (**Figure 2B**). Consistent with the role of U2AF1 specifically at the 3’SS, alternative 5’ SS (A5SS) was the least represented splicing event. Selected differentially spliced events were confirmed by splicing-specific RT-PCR validating the experimental system^34^ (**Figure S2B**).

Analysis of 3’SS sequence motifs extended observations previously reported^21–23^ (**Figure 2C**). In S34F mutant cells, in position −3 of the 3’SS region, we found higher frequency of C for more included exons (71.6%) and higher frequency of U for less included exons (85.2%). In Q157R mutant cells, more exon inclusion was associated with higher frequency of G in position +1 (65.0%) and less exon inclusion with higher frequency of A (74.7%). With respect to previous studies, our analysis revealed the same sequence-specificity also when considering S34F and Q157R differential splicing events belonging to the A3SS, MXE and RI categories (**Figures S2C-S2D**).

Importantly, these splicing-predictive positions at the 3’SS, −3 for U2AF1 S34F and +1 for U2AF1 Q157R, perfectly matched the positions where we observed *de novo* binding peaks in U2AF1 S34F and Q157R freCLIP-seq metaprofiles (**Figure 2A**). These results, based on the analysis of ~40,000 intron-exon junctions *in vivo* at single-nucleotide resolution, suggest a direct relationship between changes in RNA binding and splicing outcomes.

### Integrative binding-splicing analysis reveals that U2AF1 mutant gain-of-binding can lead to loss-of-splicing

Transcripts that are differentially bound and spliced are likely directly affected by splicing factor mutations. To better understand how RNA binding changes driven by U2AF1 mutations alter RNA splicing in myeloid malignancies, we performed an integrative multi-omics analysis identifying intron-exon junctions affected by both differential binding (freCLIP-seq) and splicing (RNA-seq) in U2AF1 S34F and Q157R cells. Differentially bound junctions were identified comparing both light and heavy fractions in freCLIP-seq (see Methods; **Figures S3A-S3B**, **Table S3**).

Taking into consideration all the possible combinations of increased *vs* decreased mutant binding and increased *vs* decreased junction inclusion, we analyzed four binding-splicing classes (see Methods; **Table S4**). The “<binding;<inclusion” and “>binding;>inclusion” classes, with congruent binding-splicing outcomes, match the previously proposed model^21,23,24^ where more or less mutant U2AF1 binding directly results into more or less exon inclusion. Conversely, the “>binding;<inclusion” and “<binding;>inclusion” classes suggest an “inverse” model, where mutant U2AF1 binding is inversely related to exon inclusion. In this scenario, increased mutant U2AF1 binding can result in reduced exon inclusion, possibly mediated by impaired splicing progression.

Focusing on the most highly represented differential splicing event, skipped exons (**Figure 2B**), we identified several junctions confirming the previously proposed model for the S34F mutant: “<binding;<inclusion”, 67 junctions; “>binding;>inclusion”, 71 junctions (**Figure 3A**, central panel). However, 55.3% of the aberrantly bound and spliced junctions followed the “inverse” model with “>binding;<inclusion” as the most enriched class (123 out of 309 junctions). Metaprofiles based on the four binding-splicing classes showed that increased U2AF1 S34F binding in position −3 of the 3’SS was associated with a reduction in U2AF2 binding, particularly evident for skipped exon junctions (**Figure 3A**, top panels). On the contrary, a decrease in U2AF1 S34F binding was accompanied by increased U2AF2 binding, especially for more included exons (**Figure 3A**, bottom panels). Quantitation of binding differences (delta analysis) confirmed these trends, suggesting that more *vs* less mutant U2AF1 binding resulted in splicing alterations by affecting U2AF2-RNA interactions in the opposite direction (**Figures S4A-S4B**).

**Figure 3.**
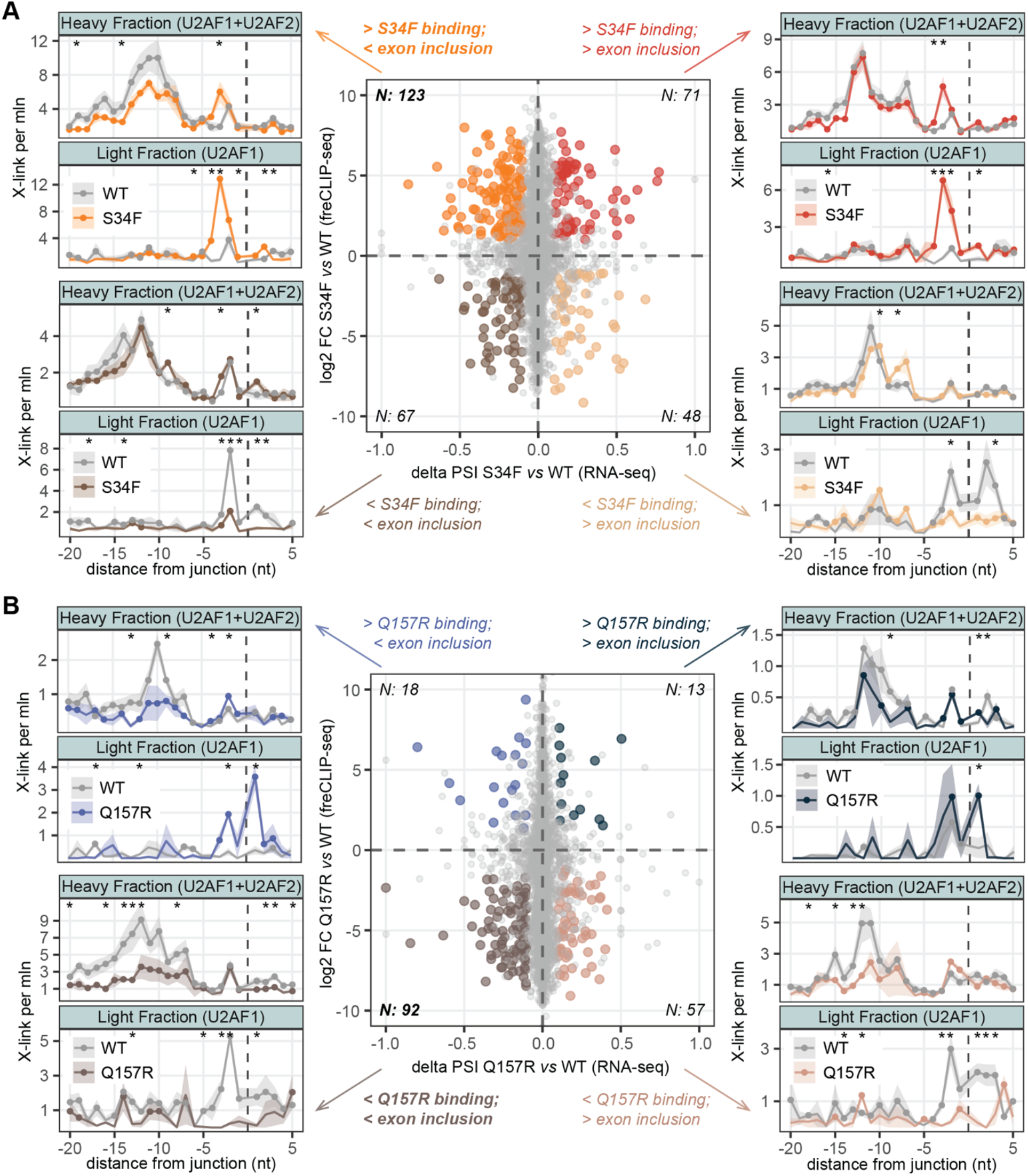
Multifaceted relationship between differential binding-splicing in U2AF1 mutants. **(A, B)** Central panel: scatter plot representing intron-exon junctions significantly affected by both differential binding (y-axis, log2 FC based on freCLIP-seq) and differential splicing (x-axis, delta PSI of SE events based on RNA-seq) in S34F *vs* WT **(A)** and Q157R *vs* WT **(B)**. Significantly affected junctions (combined binding-splicing P-value<0.05, Fisher’s method) in each binding-splicing class (“<binding;<inclusion”, “>binding;>inclusion”, “>binding;<inclusion”, “<binding;>inclusion”) are color-coded. N, number of significantly affected junctions in each class. Side panels: U2AF binding metaprofiles (mean±SEM of the number of crosslinking events per million reads), based on heavy and light freCLIP-seq fractions, considering junctions belonging to the four binding-splicing classes in S34F *vs* WT **(A)** and Q157R *vs* WT **(B)**. Positions characterized by a significant change in mutant vs WT U2AF1 binding are starred (P-value<0.05, one-tailed t-test).

The same analysis for the Q157R mutant revealed a different scenario with fewer affected junctions than for S34F (180 *vs* 309), most of which exhibited a loss-of-binding pattern: 51.1% of junctions in the “<binding;<inclusion” class and 31.7% of junctions in the “<binding;>inclusion” class (**Figure 3B**, central panel). Metaprofiles based on the four binding-splicing classes showed a general reduction in U2AF2 signal, with prominent binding of Q157R mutant at position +1 of the 3’SS (**Figure 3B**, side panels; **Figures S4C-S4D**).

Collectively, in-depth integrative analysis of *in vivo* binding and splicing alterations revealed a complex picture that expands upon previous models inferred from splicing and *in vitro* binding data. Mutation-induced gain-of-binding may result in exon skipping, possibly mediated by compromised recruitment of U2AF2.

### U2AF1 S34F and Q157R mutations converge on key biological processes

Hotspot S34 and Q157 mutations occur in the two zinc fingers of U2AF1 that are critical to RNA binding^5^. Understanding common and distinct splicing and binding perturbations associated with these two mutations can shed light on their biological implications in cancer.

We conducted a comprehensive meta-analysis of 18 published datasets^21–24,35–41^ cataloging aberrant splicing patterns secondary to pathogenic U2AF1 mutations in our HEL cell model and in published engineered cell lines, mouse models and primary samples (**Table S5**). The analysis of 17 S34F/Y datasets revealed a total of 3,740 differentially spliced genes identified in only one dataset, suggesting that splicing changes can be cell-system dependent, as previously reported^22,42^ (**Figures 4A-4B**, **Table S6**). Splicing alterations in 880 genes were uniquely present in our system, whereas 2,726 genes were identified as aberrantly spliced also in other datasets (**Figure 4B**, **Table S6**). Similar conclusions can be drawn from the analysis of Q157R/P datasets, although this mutant causes fewer splicing alterations and only 2 out of 18 datasets were available^21,40^ (**Figures 4C-4D**, **Table S6**). Importantly, in our system we detected more than half of the genes (54.6%) reported as aberrantly spliced in seven patient-based datasets (**Figures 4A-4C**, **Table S6**).

**Figure 4.**
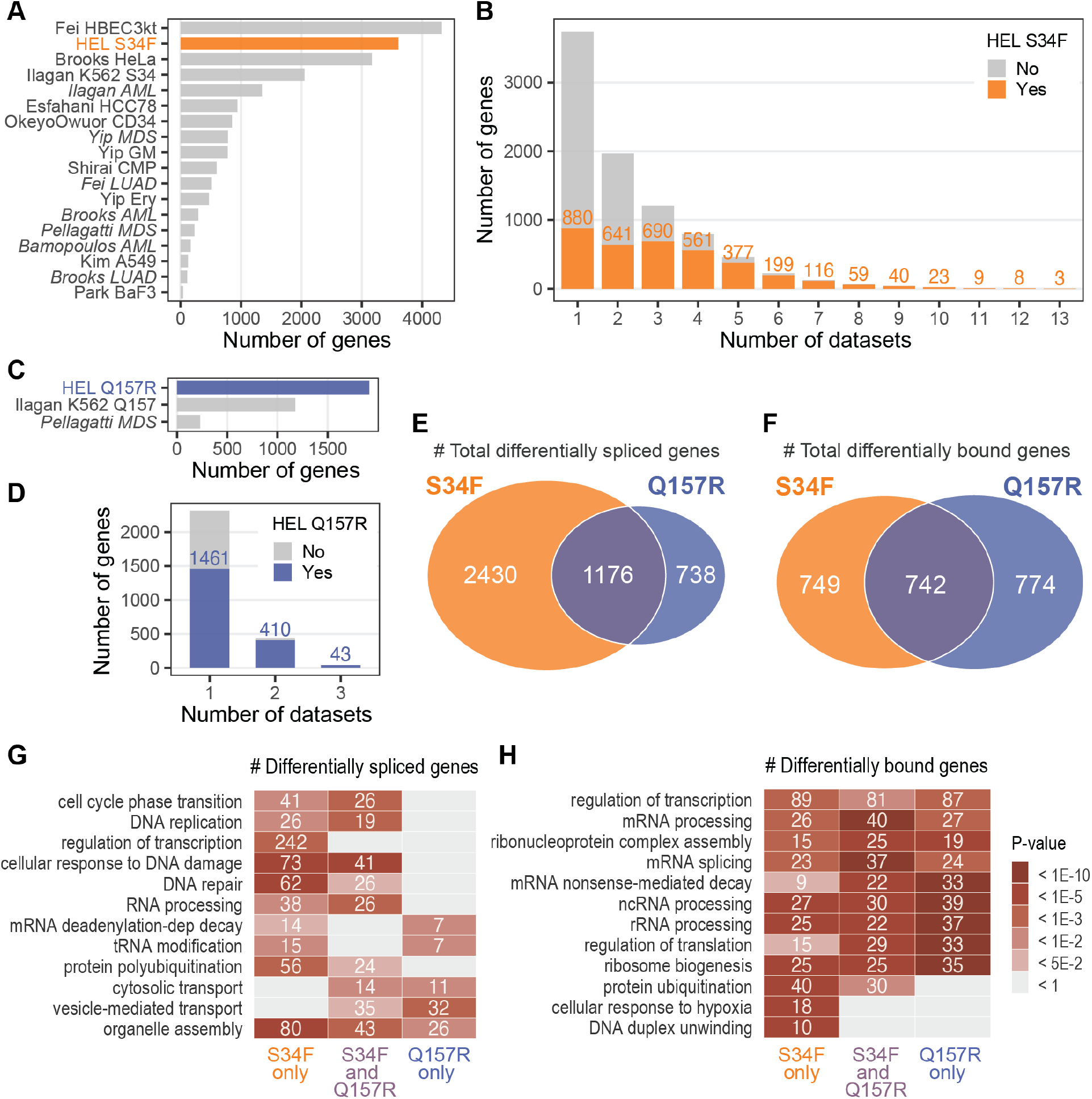
Comprehensive analysis of splicing and binding alterations reveals convergent pathways impacted by U2AF1 mutations. **(A, C)** Number of differentially spliced genes in U2AF1 S34F/Y (**A**) or Q157R/P (**C**) condition identified in our HEL system and in the meta-analysis of published datasets (**Table S3**). Patient-based datasets are indicated in italic. **(B, D)** Histogram showing the overlap between S34F/Y (**B**) or Q157R/P (**D**) differentially spliced genes across RNA-seq datasets. For each bar, the number of genes differentially spliced in our HEL dataset is highlighted. **(E)** Intersection between differentially spliced genes (absolute delta PSI>10%; FDR<0.05) in S34F *vs* WT and Q157R *vs* WT HEL cells. **(F)** Intersection between differentially bound genes (absolute log2 FC>0.75; P-value<0.05) in S34F *vs* WT and Q157R *vs* WT HEL cells, determined by freCLIP-seq. **(G)** Gene Ontology enrichment analysis (Biological Process) of differentially spliced (left panel) and differentially bound (right panel) genes in S34F, Q157R or both mutant conditions. Heatmaps are colored according to the significance of the enrichments and contain the number of genes falling into each category.

While splicing alterations were the predominant events, U2AF1 mutations also caused changes in gene expression: we identified 1,129 differentially expressed genes in S34F mutant cells (**Figure S5A**) and 98 differentially expressed genes in Q157R mutant cells (**Figure S5B**).

Comparing differential splicing events among the two mutants, we obtained 2,430 genes differentially spliced exclusively in S34F, 738 exclusively in Q157R and 1,176 in both U2AF1 mutants (27.1% of overlap) (**Figure 4E**). Considering differential binding, we identified 749 genes affected exclusively in S34F cells, 774 genes exclusively in Q157R cells, and 742 in both mutants (32.8% of overlap, **Figure 4F**).

The convergence of U2AF1 mutant-induced alterations increased when we considered the gene ontology (GO) biological processes in which perturbed genes were involved. In fact, differentially spliced genes in both mutants were significantly associated with cell cycle, transcription, DNA repair, RNA processing, vesicle-mediated transport and organelle assembly (**Figure 4G**, **Table S7**); differentially bound genes in both mutants were commonly enriched in transcription, RNA processing, ribonucleoprotein complex assembly, nonsense-mediated decay and translation (**Figure 4H**, **Table S8**).

Overall, we identified a majority of distinct binding and splicing alterations induced by S34F and Q157R mutants, with a convergence on biological processes crucial to tumorigenesis.

### U2AF1 mutations alter stress granule biology

To identify biological processes directly affected by U2AF1 mutations, we performed functional analysis on all genes characterized by concurrent aberrant binding and splicing (S34F: 497 genes; Q157R: 312 genes; **Table S4**). For both S34F and Q157R mutants, we observed a significant enrichment in several GO terms related to RNA biology, in particular RNA granules: RNA transport, ribonucleoprotein complex assembly, RNA helicases, RNA binding, cytoplasmic stress granules^43,44^ (**Figure 5A**; **Table S9**). This observation was further supported by a significant enrichment in genes coding for proteins containing low-complexity domains (**Figure S6A**). These proteins promote the formation of membrane-less organelles, dynamic condensates of RNA and RNA-binding proteins, such as nucleoli and Cajal bodies in the nucleus, P-bodies and stress granules in the cytoplasm^45,46^.

**Figure 5.**
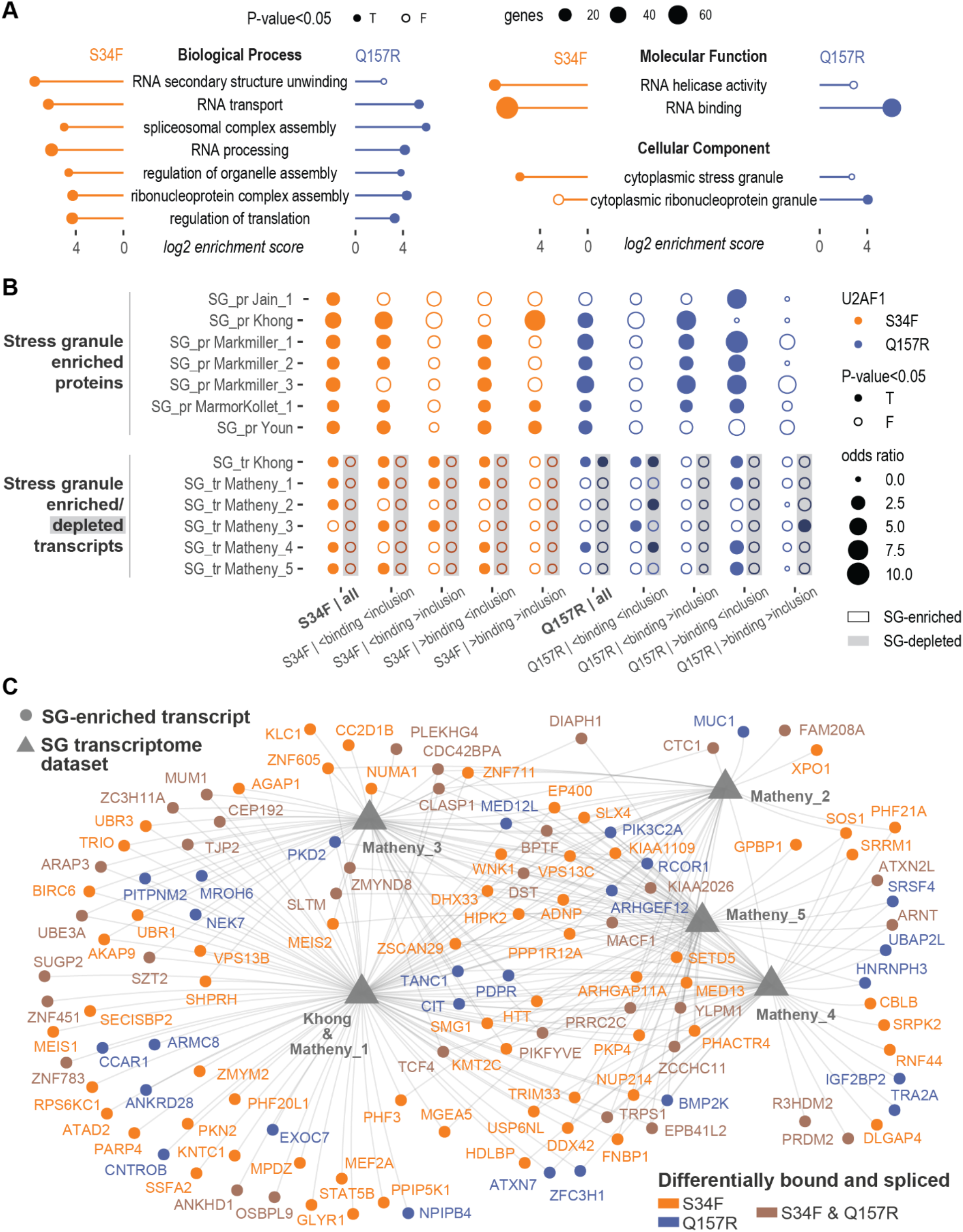
Binding-splicing alterations induced by U2AF1 mutations affect transcripts enriched in stress granules. **(A)** Enrichment in GO terms related to RNA granule biology for genes differentially bound and spliced in U2AF1 S34F *vs* WT and Q157R *vs* WT HEL cells. Node size: number of affected genes belonging to a specific term (P-value based on Fisher’s exact test). **(B)** Enrichment analysis of mutant U2AF1 differentially bound-spliced genes among SG-related experimental datasets. Top panel: SG-enriched proteins; Bottom panel: SG-enriched (white-highlighted) and SG-depleted (grey-highlighted) transcripts. Dot size is proportional to the overlap, measured by odds ratio (Fisher’s exact test). **(C)** Network visualization of differentially bound-spliced transcripts in U2AF1 mutants that are also enriched in stress granules, according to multiple SG experimental datasets.

To probe the significance of these findings, we collected previously published experimental datasets characterizing both proteins and transcripts enriched in SGs. We considered a total of 16 datasets, 10 of which identified SG-enriched proteins by mass-spectrometry and 6 of which identified SG-enriched and SG-depleted transcripts by RNA-seq^47–52^ (**Table S10**, **Figure S6B**). Importantly, we found a consistent and significant overlap between genes differentially bound and spliced in U2AF1 mutant cells and SG-enriched, but not SG-depleted, proteins and transcripts (**Figure 5B**). In particular, the “>binding;<inclusion” class was the most represented in SG-enriched transcripts and proteins for both U2AF1 mutants (**Figure 5B**). In total, 125 differentially bound-spliced genes were identified as SG-enriched transcripts (network representation in **Figure 5C**). These transcripts are characterized by long coding sequences (CDS) and untranslated regions (UTR)^47^. In addition, 97 differentially bound and spliced genes code for proteins enriched in SGs according to proteome studies (network representation in **Figure S6C**).

For 36 genes, both the transcript and the corresponding protein were enriched in SGs, such as DEAH-Box Helicase 33 (DHX33) and Kinesin Light Chain 1 (KLC1) for S34F, Ubiquitin-associated Protein 2-like (UBAP2L) and Heterogeneous Nuclear Ribonucleoprotein H3 (HNRNPH3) for Q157R, and Ataxin 2 Like (ATXN2L) and Proline Rich Coiled-Coil 2C (PRRC2C) for both mutants. To confirm alterations in SG biology in mutant U2AF1 HEL cells, we evaluated SGs at steady state and after stress induction with sodium arsenite by immunofluorescence staining against the SG-marker protein G3BP1^53^ (**Figure 6A**). Already at the steady state and especially after exposure to oxidative stress, we detected a more marked punctate G3BP1 signal in U2AF1 S34F and Q157R cells in comparison to WT cells (**Figure 6A**, **Files S1-S2**). Quantitation of SGs with an IMARIS-based image analysis pipeline, developed for accurate SG identification (**Figure S7, Table S11**), confirmed significantly higher intensity of G3BP1-marked SGs in arsenite-treated mutant compared to WT cells.

**Figure 6.**
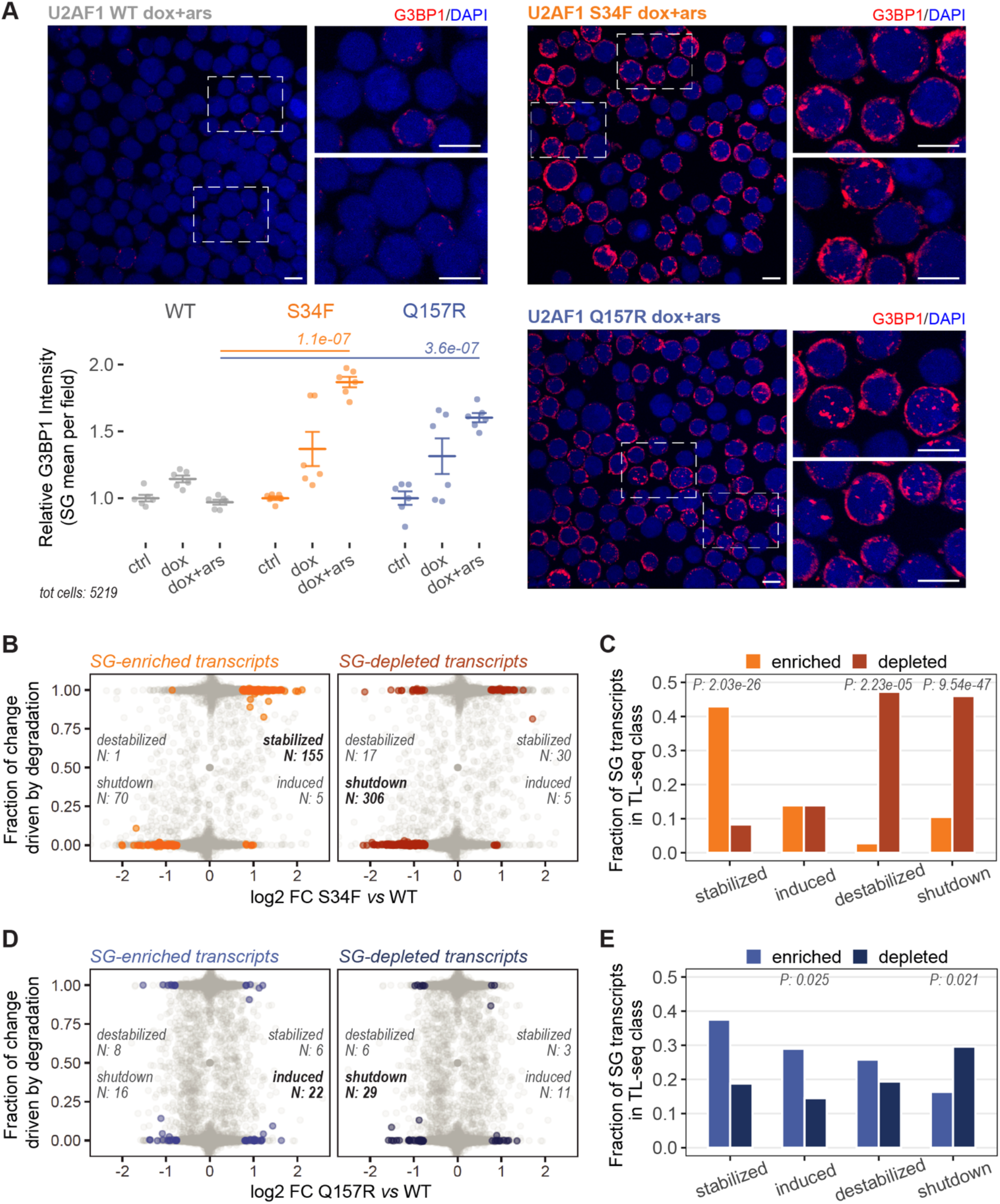
Mutant U2AF1 cells have an increased capability to form stress granules, supported by changes in RNA dynamics. **(A)** Representative IF images (scale bars, 10 μm) and quantification of stress granules in mutant and WT U2AF1 HEL cells. SGs were identified by IMARIS (see Methods). The plot displays the mean±SEM G3BP1 field intensity, normalized to the relative controls (ctrl, uninduced HEL cells without arsenite treatment; dox, doxycycline-induced HEL cells without arsenite treatment; dox+ars, doxycycline-induced HEL cells treated with 500 μM arsenite for 1 hour). G3BP1 field intensity is the mean intensity of all the single SGs identified in the field. For each condition, 6 fields were acquired (3 fields per replicate), containing on average 74 cells each. Differences between S34F or Q157R and WT were tested with two-tailed t-test. **(B, D)** Scatter plot of gene expression changes (x-axis) and relative stability/degradation contributions (y-axis) in S34F **(B)** or Q157R **(D)** vs WT, measured by TL-seq (2 replicates per condition). Transcripts enriched (left panel) or depleted (right panel) in stress granules are highlighted. N, number of transcripts in each TL-seq class (stabilized, induced, destabilized, shutdown) **(C, E)** Fraction of SG enriched *vs* depleted transcripts in each TL-seq class in S34F **(C)** or Q157R **(E)** vs WT. Differences in fractions within each class were tested with a proportion test.

To further understand mutant U2AF1-mediated pathogenic mechanisms increasing SG formation, we analyzed RNA transcript dynamics by TimeLapse-seq^54^. The TimeLapse chemistry (see Methods) allows to disentangle the contributions of RNA synthesis *versus* stability/degradation on total RNA levels. Comparing mutant *vs* WT U2AF1 cells with TL-seq, transcripts were sorted into four classes: upregulated transcripts stabilized or induced secondary to increased stability or synthesis rate, respectively; downregulated transcripts destabilized or shutdown secondary to reduced stability or synthesis rate, respectively (**Figure 6B**). By integrating TL-seq variations with the afore mentioned experimental datasets characterizing SG-enriched and SG-depleted transcripts (**Table S10**), we observed significant over-representation of SG-enriched RNAs among transcripts with increased stability (S34F, **Figures 6B-6C**) and synthesis (Q157R, **Figures 6D-6E**). Conversely, SG-depleted transcripts were mainly in the degradation/shutdown classes for both mutants (**Figures 6B-6D**). These data suggest that U2AF1 mutations increase the availability of RNAs that are prone to participate in stress granule formation.

Collectively, our molecular and imaging results support a model whereby U2AF1 mutations increase the cell’s potential to form stress granules upon stress.

### Increased stress granule formation in U2AF1-mutant primary AML samples

To explore if SG perturbations detected in our MDS/AML cell line model were also present in primary patient-derived samples, we performed G3BP1-IF staining on bone marrow or peripheral blood mononuclear cells of three AML patients with U2AF1 S34 mutation. Samples from three AML patients without U2AF1 mutations served as controls (**Table S12**). Consistent with our results in HEL cells, we observed increased formation of stress granules in U2AF1-mutant *vs* WT AML blasts upon oxidative stress (**Figure 7A-7B**, **Table S13**).

**Figure 7.**
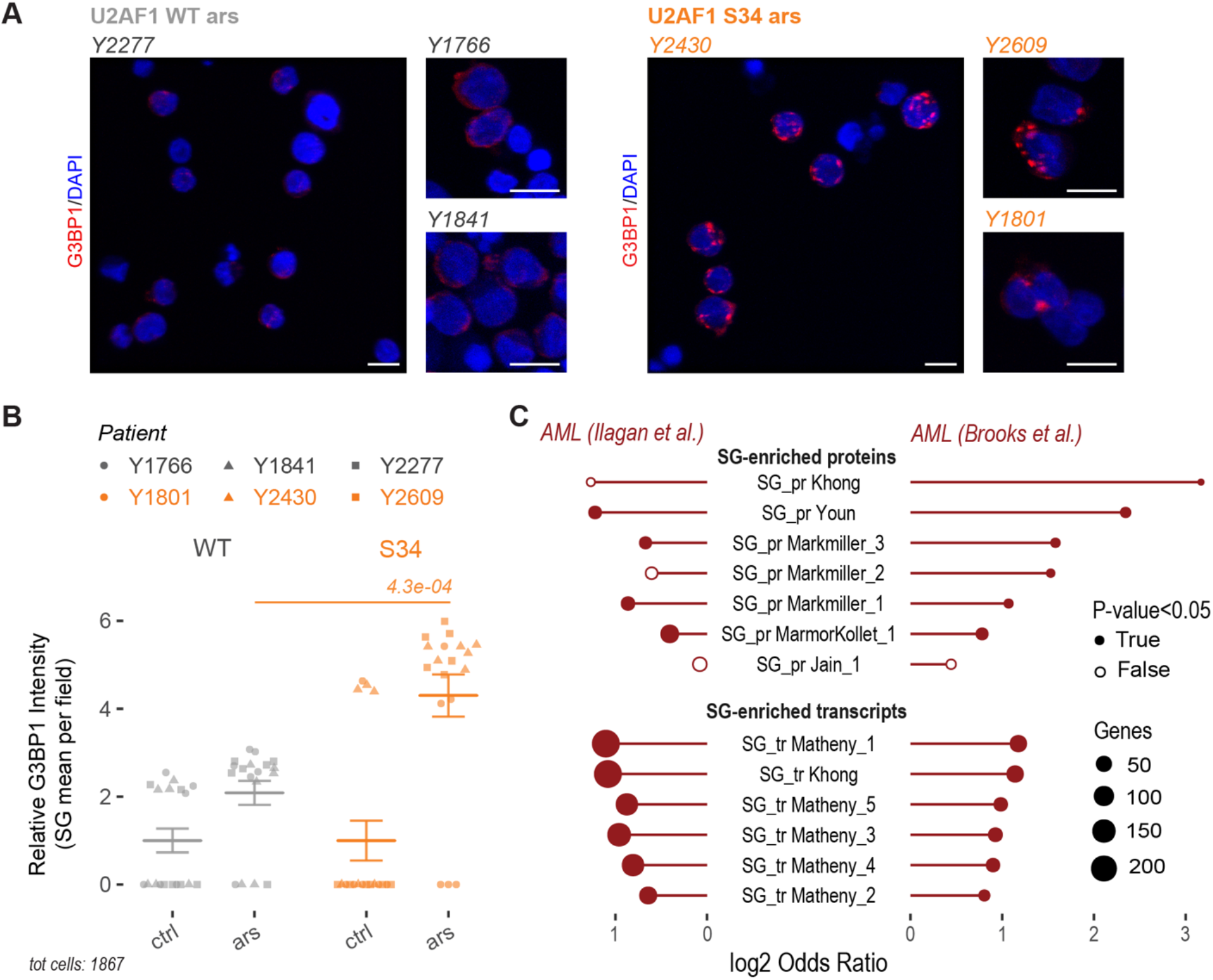
U2AF1-mutant AML patients display increased SG levels. **(A, B)** Representative IF images (scale bars, 10 μm) and quantification of stress granules in AML patients with WT (n=3) or S34F/Y (n=3) U2AF1. SGs were identified by IMARIS (see Methods). The plot displays the mean±SEM G3BP1 field intensity, normalized to the relative controls (ctrl, primary cells without arsenite treatment; ars, primary cells treated with 500 μM arsenite for 1 hour). G3BP1 field intensity is the mean intensity of all the single SGs identified in the field. For each patient, 6 fields per condition were acquired, containing on average 25 cells each. Differences between S34F/Y and WT patients were tested with two-tailed t-test. **(C)** Enrichment analysis of genes differentially spliced in 2 published cohorts of U2AF1-mutant AML patients among SG experimental datasets (Fisher’s exact test).

Differentially spliced genes from two previously published datasets on S34 *vs* WT AML patients (**Table S5**) confirmed a significant over-representation of SG-enriched proteins and SG-enriched transcripts, based on our meta-analysis of experimental SG datasets (**Figure 7C**).

This analysis, consistent with our IF imaging results, revealed an increased capability of mutant U2AF1 cells to respond to stress by forming stress granules in AML primary samples.

## Discussion

Missense mutations at codons 34 and 157 of the splicing factor U2AF1 affect 3’SS recognition leading to ineffective hematopoiesis in MDS and AML. However, critical questions remain open as to how these mutations alter *in vivo* RNA binding, splicing and kinetics and which biological processes are directly influenced by binding-splicing perturbations, contributing to the disease process. Applying a multi-omics approach on U2AF1 WT, S34F and Q157R cells we were able to: i) separate *in vivo* individual RNA binding signals of U2AF1 and U2AF2; ii) identify distinct alterations in mutant U2AF1 RNA binding, suggesting conformational changes in the U2AF1-U2AF2-RNA complex; iii) dissect complexities in the binding-splicing relationship where splicing alterations are driven not only by loss, but also by gain of mutant U2AF1 binding; iv) demonstrate that U2AF1 mutations lead to alterations in RNA granule biology, in particular stress granules, also in AML primary cells.

The application of the high-sensitivity eCLIP-seq protocol combined with the fractionation step in the U2AF1 freCLIP-seq led to a high-resolution positional analysis that, in comparison with previous U2AF1 CLIP-seq studies^32,41^, allowed to determine U2AF1- and U2AF2-specific binding contributions in the WT and mutant context. U2AF1 S34F and Q157R freCLIP-seq revealed mutant-specific changes in U2AF1 binding, with *de novo* peaks in position −3 for the S34F mutant and in position +1 for the Q157R mutant, together with a general reduction in U2AF2 signal. Crucially, these peaks perfectly matched sequence-specific positions affected by aberrant splicing in RNA-seq data. We validated our set of differentially spliced genes against 18 published datasets^21–24,35–41^ and in addition were able to fully characterize sequence-specific differences also for less frequent events such as alternative 3’ splice sites and intron retention.

Preliminary studies suggested that S34 and Q157 mutations determine distinct molecular and clinical features in myeloid malignancies and other cancers^55,56^. We indeed observed distinct splicing and binding alterations in S34F and Q157R mutants at the junction level, however, pathway enrichment analysis pointed towards convergence on critical biological processes relevant in tumorigenesis.

The literature to date suggests that aberrant splicing directly results from loss of mutant U2AF1 binding at specific junctions^21,23,24^. A recent study, assessing *in vitro* U2AF1 binding, questioned sequence-specific reductions in binding; U2AF1 S34F compared to WT showed little discrimination for the −3 nucleotide sequence, with similar affinity to any nucleotide except G^25^. Similarly, limited −3 sequence specificity was also observed in studies of the effects of S34F mutation on the open *vs* closed conformations of U2AF2^57^. Our integrative analysis unveiled an unexpectedly multifaceted relationship between mutant U2AF1 binding and splicing outcomes. While Q157R mainly induced a loss-of-binding pattern where mutant binding and splicing were directly proportional, S34F predominantly supported a gain-of-binding process, where increased binding instead led to reduced U2AF2 binding and reduced exon inclusion.

Our in-depth integrative binding-splicing analysis also offered the possibility to discover novel biological processes directly influenced by pathogenic U2AF1 mutations in the context of myeloid malignancies. We specifically noticed, among perturbed genes, a significant enrichment in stress granule-related processes. SGs are membrane-less organelles constituted by multiple RNAs and RNA binding proteins. They are formed, prevalently upon stress, in the cytoplasm of eukaryotic cells, improving cellular adaptation to stress conditions^58^. Increased SG formation has been linked to tumorigenesis as a strategy exploited by cancer cells to enhance their fitness under stress^59–61^. We confirmed the enrichment of differentially bound-spliced genes in a collection of 16 experimental datasets characterizing SG-enriched proteins and transcripts by mass spectrometry and RNA-seq respectively. By IF imaging, we revealed a marked increase in stress granules in U2AF1 S34F and Q157R over WT cells upon arsenite treatment, indicating an increased capability of mutant cells to aggregate RNA-protein granules when facing stress. Moreover, analysis of RNA dynamics confirmed that U2AF1 mutations increased the stability/synthesis of transcripts enriched in SGs, and conversely promoted the degradation/shutdown of transcripts depleted in SGs, providing a molecular explanation for the increase in SG observed by imaging. Importantly, our observations on the increase in stress granules were translationally validated through IF analysis of U2AF1-mutant *vs* WT primary AML samples. These results lay the foundation for a new paradigm where mutations in splicing factors ultimately alter membrane-less organelles by acting on the availability of their RNA and protein components (**Figure S8**).

From a therapeutic perspective, current clinical approaches for targeting RNA splicing include antisense oligonucleotides, inducing splice-site switches of specific RNA isoforms, and small-molecule modulators directed against spliceosomal components^62,63^. In a gain-of-binding but not in a loss-of-binding model, anti-sense oligonucleotides may be of utility. Moreover, the gain-of-binding/loss-of-splicing mechanism could be exploited to exacerbate splicing defects in U2AF1 mutant cells. Indeed, Chatrikhi *et al.*^70^ recently described a compound that inhibits *in vitro* splicing by stalling the spliceosome machinery through increased U2AF2-RNA binding.

Splicing factor-mutant hematopoietic cells rely on the wild-type counterpart for their survival, rendering them sensitive to pharmacologic inhibition of splicing catalysis^24,64^. To date, the full efficacy of spliceosome inhibitors in the treatment of U2AF1-mutant myeloid malignancies remains to be shown^65–67^. Protein arginine methyltransferase (PRMT) inhibitors, that exhibit a strong anti-leukemic effect on spliceosomal mutant AML in preclinical studies^67^, induce methylation changes in numerous intrinsically disordered proteins, such as G3BP1, suggesting an impact on membrane-less organelle formation. Our results highlight the relevance of future studies to: i) further characterize SG perturbations in the presence of splicing factor mutations; ii) assess the therapeutic potential of drug combinations targeting stress granules and spliceosome. Since SG components contribute to various cancer-related processes such as cell cycle progression, apoptosis inhibition, resistance to stress and therapeutics, targeting SGs may represent an effective therapeutic approach^68,69^.

Collectively, our findings provide detailed insights into U2AF1 mutation-dependent pathogenic RNA mechanisms that will be useful in the development of targeted therapeutics against U2AF1-mutant myeloid malignancies and other cancers.

## Methods

### Cell lines

All cell lines were cultured under 5% CO_2_ at 37°C. 293FT cells for lentivirus production were grown in DMEM (ThermoFisher SCIENTIFIC, Cat #11965092) supplemented with 10% FBS (Gemini Bio-products, Cat #100-106). HEL erythroleukemia cells (ATCC, Cat #TIB-180) were grown in RPMI 1640 (ThermoFisher SCIENTIFIC, Cat #11875093) supplemented with 10% FBS (Gemini Bio-products, Cat #100-106) before transduction or 9% tetracycline-negative FBS (Gemini Bio-products, Cat #100-800) after transduction, 1% L-glutamine (ThermoFisher SCIENTIFIC, Cat #25030081) and 1% penicillin-streptomycin (pen-strep; ThermoFisher SCIENTIFIC, Cat #15140122).

### Human subjects

Human primary cells were obtained with patients’ written consent after approval by the Yale University Human Investigation Committee. We included in our cohort 6 patients (**Table S12**) affected by AML and characterized by U2AF1 WT (n=3) or S34F/Y (n=3), as reported by next-generation sequencing report using Yale 49-gene myeloid panel (Ion Torrent platform). Mononuclear cells were isolated by density gradient centrifugation from BM or PB samples collected at the time of diagnosis and frozen in FBS+10% dimethyl sulfoxide. After thawing, primary cells were cultured overnight in StemSpan (STEMCELL Technologies, 09650) supplemented with 1% pen-strep and recombinant human cytokines FLT-3 (50 ng/mL), SCF (50 ng/mL), TPO (100 ng/mL), IL-3 (10 ng/mL) and IL-6 (25 ng/mL). All cytokines were purchased from Gemini Bio-products.

### U2AF1 cell line generation and verification

Full length human WT, S34F or Q157R FLAG-tagged U2AF1 in CS-TRE-Ubc-tTA-I2G plasmids (**Figure 1B**), encoding for tetracycline-responsive element (TRE) and enhanced green fluorescent protein (EGFP), were a kind gift from Tomoyuki Yamaguchi at Japan Science and Technology Agency^5,71^. Lentivirus production was obtained by co-transfecting 293FT cell line with psPAX2 (Addgene, plasmid #12260), pCMV-VSVG (Addgene, plasmid #14888) and U2AF1-containing plasmid, followed by spin-concentration. HEL cells were infected with viral supernatants via spinoculation (1000 g for 90 min at 30°C) with addition of 4 μg/mL polybrene (Sigma-Aldrich, Cat #H9268). 48 hours after transduction, GFP^+^ cells (mean: WT = 21.2%, S34F = 22.2%, Q157R = 27.4%) were sorted by fluorescence-activated cell sorting (FACSAria II, BD Biosciences, Yale Flow Cytometry Facility). To express FLAG-tagged U2AF1 proteins, HEL cells were induced with 1 μg/mL doxycycline for 48 hours and the expression was verified through PCR followed by Sanger sequencing (3730xL DNA Analyzer, ThermoFisher SCIENTIFIC, Yale Keck DNA Sequencing Facility) and through western blotting (**Figure S1A**).

### RNA extraction, reverse-transcription and cDNA amplification

RNA was isolated using the RNeasy Mini kit (QIAGEN, Cat #74104) following manufacturer’s instructions. One microgram of extracted RNA was reverse transcribed into cDNA using iScript cDNA Synthesis Kit (BIO-RAD, Cat #1708890). Amplification of cDNA for validation of doxycycline induction was performed with U2AF1 primers (forward: 5’-GGCACCGAGAAAGACAAAGT-3’; reverse: 5’-CTCTGGAAATGGGCTTCAAA-3’) and PCR products were purified using QIAquick PCR purification kit (QIAGEN, Cat #28104) before Sanger sequencing. Three representative alternative splicing events were validated, in triplicate, with previously reported target specific primers^34^. PCR products were resolved by agarose gel electrophoresis, visualized using Image Lab 3.0 software (BIO-RAD), and quantified in ImageJ.

### Western blotting

Cellular lysates were incubated at 95°C for 5 min and separated in 12% Mini-PROTEAN TGX Precast Protein gels (BIO-RAD, Cat #4561044). Proteins were then transferred to 0.45 μm PVDF membrane at 100V for 1 hr. Membranes were blocked with 5% skim milk for 40 min, incubated with primary antibodies overnight at 4°C and incubated with secondary antibodies for 1 h at room temperature. Washing steps were performed using 1X TBST. Membranes were developed with SuperSignal West Femto Maximum Sensitivity Substrate (ThermoFisher SCIENTIFIC, Cat #34095). Antibodies were used at the following dilutions: rabbit polyclonal anti-U2AF1 (Bethyl Laboratories, Cat #A302-079) 1:5000, mouse monoclonal anti-FLAG M2 (Sigma-Aldrich, Cat #F1804) 1:1000, rabbit monoclonal anti-G3BP1 (Abcam, Cat #ab181149) 1:5000, mouse monoclonal anti-HSP90 (StressMarq Biosciences, Cat #SMC-107B) 1:5000, rabbit polyclonal anti-GAPDH (FL-335, Santa Cruz Biotechnology, Cat #sc-25778) 1:5000, goat anti-rabbit IgG HRP-linked (Cell Signaling TECHNOLOGY, Cat #7074) 1:5000, horse anti-mouse IgG HRP-linked (Cell Signaling TECHNOLOGY, Cat #7076) 1:5000. Chemiluminescence was visualized by Image Lab 3.0 and protein bands’ quantification was performed in ImageJ.

### eCLIP-seq

eCLIP-seq experiments (U2AF1 eCLIP-seq, U2AF2 eCLIP-seq, U2AF1 freCLIP-seq) were performed at least in duplicate (**Table S1**), as per ENCODE guidelines (https://www.encodeproject.org), according to the published protocol^26^ with the following modifications: HEL cells containing FLAG-tagged U2AF1 WT, S34F or Q157R were treated with doxycycline (1 μg/mL) for 48 hours before UV-crosslinking (400 mJ/cm^2^ UV Stratalinker 2400, STRATAGENE). RNA-protein complexes were immunoprecipitated with 12 μg anti-FLAG M2 antibody (Sigma-Aldrich, Cat #F1804) or 8 μg anti-U2AF2 antibody (Sigma-Aldrich, Cat #U4758) and Dynabeads Protein G (ThermoFisher SCIENTIFIC, Cat #10004D). RNA was partially digested with RNase I (ThermoFisher SCIENTIFIC, Cat #AM2295) and P32-labeled (PerkinElmer, Cat #BLU002Z250UC), followed by RNA linker ligation as per the relevant protocol. U2AF-RNA complexes were isolated by SDS-PAGE and transferred to nitrocellulose membranes. For U2AF1 freCLIP, membrane region between 37-65 kD was excised to obtain the “light fraction” containing U2AF1 monomer plus bound RNA, and region between 65-110 kD was excised to obtain the “heavy fraction” containing the U2AF heterodimer plus bound RNA (**Figure S1C**). Light and heavy fractions were obtained for each sample. For both eCLIP-seq and freCLIP-seq, excised membranes were treated with proteinase K and the RNA was isolated. Reverse transcription and library preparation were carried out according to the published protocol. Libraries were deep-sequenced on Illumina HiSeq2500 system, paired-end 75 bp, and on Illumina HiSeq4000 system, paired-end 100bp, at the Yale Center for Genome Analysis (YCGA).

### RNA-seq

Total RNA from induced U2AF1 WT, S34F and Q157R HEL cells (in duplicate as per the ENCODE guidelines, https://www.encodeproject.org), as well as from uninduced HEL cells (in duplicate, as controls), was extracted using the RNeasy Mini kit (QIAGEN, Cat #74104) and submitted to library preparation and sequencing at the YCGA. Ribosomally depleted RNA was sequenced on Illumina HiSeq4000 system, paired-end 100bp.

### TL-seq

Doxycycline-Induced U2AF1 WT, S34F and Q157R HEL cells and not-infected HEL cells were labeled in duplicate with 100 μM of the uridine analog 4-thiouridine (s^4^U, Alfa Aesar, Cat #AAJ60679MC) for 2 hours at 37°C in the dark. Total RNA from labeled samples and from not-labeled control was isolated using 1 mL TRIzol (ThermoFisher SCIENTIFIC, Cat #15596-018), purified and treated with TL chemistry as reported^54^. In brief, RNA isolated from TRIzol was precipitated in 50% isopropanol supplemented with 1 mM DTT. Genomic DNA was depleted by treating with TURBO DNase (ThermoFisher SCIENTIFIC, Cat #AM2239) and RNA was purified with one volume of Agencourt RNAclean XP beads (Beckman Coulter, Cat #A63987) according to manufacturer's instructions. 5 μg of total RNA was subjected to TL chemistry for one hour at 45°C followed by reducing treatment for 30 minutes at 37°C. 10 ng of RNA was used to prepare cDNA sequencing libraries with the Clontech SMARTer Stranded Total RNA-Seq kit (Pico Input) with ribosomal cDNA depletion (Takara Bio USA, Cat #634413). Prepared libraries were submitted to paired-end 100 bp sequencing on a NovaSeq 6000 system at the YCGA. The TL chemistry allows to recode the hydrogen bonds of the uridine analog to match those of cytosine, thereby introducing U-to-C mutations in newly transcribed RNAs during reverse transcription. After alignment of the sequencing data, the level of T-to-C mutations is used to assess RNA synthesis (high T-to-C mutation rates) *vs* stability (low T-to-C mutation rates).

### Immunofluorescence staining and confocal microscopy

Immunofluorescence staining for stress granules’ analysis was conducted on induced and uninduced U2AF1 WT, S34F and Q157R HEL samples (in duplicate), and on AML primary samples (WT, n=3; S34F/Y, n=3). Samples treated with 500 μM sodium arsenite (Sigma-Aldrich, Cat #S7400) for 1 hour and not-treated samples were collected. After PBS wash, primary cells were fixed with 4% paraformaldehyde for 15 min at room temperature. Fixation step was followed by permeabilization with 100% methanol (pre-chilled to −20°C) for 10 min at room temperature as reported in SG staining published protocol^53^. Cells were then rinsed in PBS, blocked with 8% donkey serum (Abcam, Cat #ab7475) for 45 min at room temperature, rinsed in PBS, and incubated with primary antibody rabbit monoclonal anti-G3BP; (Abcam, Cat #ab181149) 1:300 for 1 hour at room temperature. After 3x PBS washes, cells were incubated with secondary antibody donkey anti-rabbit IgG Alexa Fluor 555 conjugate (Abcam, Cat #ab150062) 1:500 for 1 hour at room temperature. After 2x PBS washes, cells were spun onto glass slide and covered with DAPI-containing ProLong Gold Antifade Mountant (ThermoFisher SCIENTIFIC, Cat #P36935) and coverslip. Each step was followed by centrifugation at 500xg for 5 min. Z-stack images (HEL samples: zoom=1.7X; n steps=10-12, step size=0.38μm, 3 fields/slide; primary samples: zoom=2.5X; n steps=32-40, step size=0.25μm, 6 fields/slide; image format=1024×1024 pixels) were acquired by Leica TCS SP5 confocal microscope with 63X NA 1.40 oil objective.

### Data analysis

#### eCLIP-seq

eCLIP-seq reads were processed using FastUniq (http://sourceforge.net/projects/fastuniq/) for duplicate removal and Cutadapt (https://cutadapt.readthedocs.org) for adapter trimming. After quality control (FastQC, https://www.bioinformatics.babraham.ac.uk/projects/fastqc/), reads were aligned to the human genome (GRCh38.p10) with STAR (version 2.7.0f, --quantMode GeneCounts), using the GENCODE Release 27 for transcript annotation. BAM files were converted into BED files (bedtools, https://github.com/arq5x/bedtools2) to extract the genomic position of the crosslinked nucleotide right after the end of each sequenced read and were then re-converted into single nucleotide BAM files for post-processing analysis. Intron-exon junctions (interval from −40 to +10 around the 3’SS) were filtered using a coverage threshold of at least 10 reads in at least two replicates to remove junctions with low signal. Normalization was performed with the TMM method implemented in the edgeR Bioconductor package (https://bioconductor.org/packages/edgeR/). Differential analysis of mutant vs WT U2AF1 binding sites was performed comparing “light” vs “heavy” fraction signals in freCLIP-seq data. We specifically considered genomic positions covered by at least 5 reads in all replicates and in an interval from −4 to +2 around the 3’SS in the light fractions (U2AF1 signal) or from −20 to −5 in the heavy fractions (U2AF2 signal). Significant differentially bound sites were identified applying the “*glmQLFTest*” function in edgeR with the following thresholds: mean normalized counts per million (CPM) > 1 in either mutant or WT U2AF1 samples; absolute log2 FC > 0.75; P-value < 0.05. We defined events as characterized by increased mutant U2AF1 binding when the signal of the mutant over the wild-type was shifted toward the light fraction (**Figure S3C**) and, on the opposite, by decreased mutant U2AF1 binding when the signal of the mutant was shifted toward the heavy fraction (**Figure S3D**).

#### RNA-seq

RNA-seq reads were processed with FastUniq to remove duplicates and aligned to the human genome (GRCh38.p10) with STAR (version 2.7.0f, --quantMode GeneCounts), using the GENCODE Release 27 for transcript annotation. Normalization with the TMM method was performed with the edgeR package. To identify differentially expressed genes, we applied the “*glmQLFTest*” function in edgeR considering 2 factors: genotype (3 levels: WT, S34F, Q157R) and treatment (2 levels, uninduced and doxycycline-induced). Significant genes were then filtered according to the following thresholds: CPM > 1; absolute log2 FC > 0.75; P-value < 0.05. Alternative splicing analysis was performed with rMATS v4.0.2 (http://rnaseq-mats.sourceforge.net), capable of handling replicates with high processing speed. Alternative splicing events with absolute difference in percent spliced-in (delta PSI) > 10% and FDR < 0.05 were considered significant. Events in the comparison induced *vs* uninduced U2AF1 WT HEL cells (dox-dependent events) were removed from the final list of differentially spliced events in mutant U2AF1 conditions.

#### Integrative binding-splicing analysis

Differential binding and alternative splicing data were integrated by performing Fisher’s method through the metaseqR Bioconductor package (https://bioconductor.org/packages/metaseqR/). Only events with combined P-value < 0.05 were considered as significantly affected by aberrant binding and aberrant splicing. The four classes describing all the possible relationships between binding and cassette exons in mutant vs WT U2AF1 conditions (“<binding;<inclusion”, “>binding;>inclusion”, “>binding;<inclusion”, “<binding;>inclusion”) were defined considering absolute log2 FC (freCLIP-seq analysis) > 1 and absolute delta PSI (RNA-seq analysis) > 10%.

#### Functional annotation enrichment analysis

Pathway enrichment for junctions affected by differential binding (freCLIP-seq), differential splicing (RNA-seq) and differential binding-splicing (integrative analysis) was evaluated using enrichR package (https://CRAN.R-project.org/package=enrichR) considering all the available databases.

#### TL-seq

Filtering and alignment to the human GRCh38 genome version 26 (Ensembl 88) were performed essentially as previously described^54^. Briefly, reads were trimmed of adaptor sequences with Cutadapt v1.16 and aligned to the GRCh38 genome with HISAT2 with default parameters and -- mp 4,2. Reads aligning to annotated transcripts were quantified with HTSeq (https://pypi.org/project/HTSeq/) htseq-count. SAMtools v1.5 was used to collect only uniquely mapped read pairs (SAM flag = 83/163 or 99/147). For mutation calling, T-to-C mutations were not considered if the base quality score was less than 40 and the mutation was within 3 nucleotides from the read’s end. Sites of likely single-nucleotide polymorphisms (SNPs) and alignment artefacts (identified with bcftools) and sites of high mutation levels in the non-s4U treated controls (binomial likelihood of observation p < 0.05) were not considered in mutation calling. Normalization scale factors were calculated with edgeR using calcNormFactors (method = ‘upperquartile’). Browser tracks were made using STAR (version 2.5.3a) and visualized in IGV (https://software.broadinstitute.org/software/igv/).

Changes in expression between WT and S34F or Q157R were evaluated with DESeq2 (https://bioconductor.org/packages/DESeq2/), and genes with absolute log2 FC > 0.75 and P-value < 0.05 were considered significant. RNA kinetic parameters were estimated with a Bayesian hierarchical modeling approach using RStan software (version 2.19.3, https://mc-stan.org/rstan) as previously reported^54^. We extracted the 80% confidence interval of the fraction of change in total RNA attributed to degradation (*frac_deg_*) to identify genes whose change could be attributed primarily to changes in stability (*k_deg_*) or synthesis (*k_syn_*). If the 80% credible interval does not overlap 0.5, the gene is confidently driven by changes in stability (*frac_deg_* > 0.5) or synthesis (*frac_deg_* < 0.5). TL-seq classes combine expression and kinetic changes and are defined according to the following parameters: stabilized or induced genes, log2 FC > 0.75 and *frac_deg_* > 0.5 or < 0.5, respectively; destabilized or shutdown genes, log2 FC < −0.75, *frac_deg_* > 0.5 or < 0.5, respectively.

#### Immunofluorescence image analysis

Images were analyzed using IMARIS (Oxford Instruments, version 9.6). Specifically, ImarisCell was used to identify, segment, measure, and analyze cell, nucleus and vesicles (stress granules) in 3D. Nuclei were identified based on intensity, using automatic thresholding settings. Stress granules from HEL samples were identified by first using an estimated diameter of 1 μm, and then refining the selection with mean intensity and quality settings set to automatic, and intensity standard deviation setting, to select voxels above 22.0 intensity units. Stress granules from primary samples were identified by first using an estimated diameter of 0.6 μm, and then refining the selection with mean intensity and voxel number settings of 25.0 and 46.0 respectively.

#### Statistical analysis

Statistical analyses were performed in R (https://www.r-project.org). Number of replicates and performed statistical tests are defined in the figure legends. P-values < 0.05 were considered statistically significant and indicated within figure panels.

## Data availability

Sequencing files generated from this work have been deposited in the GEO database and are available under the accession number GSEXXXXX.

## Acknowledgements

We thank all our patients and all clinical staff for their help with patient recruitment. We thank Tomoyuki Yamaguchi for CS-TRE-Ubc-tTA-I2G plasmid, Didier Trono for psPAX2 plasmid (Addgene, #12260) and Tannishtha Reya for pCMV-VSVG plasmid (Addgene, #14888). We thank Christopher Castaldi and the YCGA for high-throughput sequencing, Yale Center for Research Computing for the use of clusters, Thomas Ardito for Leica TCS SP5 training, Jane Huang for helping in RNA extraction, Martina Cusan and Wei Liu for IF advices, Diane Krause, Clara Kielkopf and Manoj Pillai for helpful suggestions. This study was funded in part by Edward P. Evans Foundation, by the NIH/NIDDK R01DK102792, YCCC pilot grant (to SH) and CT STEM. TT was supported by a pilot grant from the Yale Cooperative Center of Excellence in Hematology (YCCEH) (NIDDK U54DK106857) and by AIRC under MFAG 2020 - ID. 24883 project. YS was supported by the Young Scientists Fund of the National Natural Science Foundation of China (grant n. 81800122). GV was supported by CNR Short Term Mobility 2018. MDS was supported by NIGMS 5R01GM137117.

## Author contributions

Conceptualization: GB, PJ, TT and SH; Methodology: GB, PJ, HL, JB, MDS, KMN, TT and SH; Investigation: GB, PJ, TH, JTZ and YG; Data Analysis: GB, PJ, TH, JTZ, YG, MDL, EC, AESB, AQ; Bioinformatics: GB, JTZ, MDL, MM and TT; Validation: VB, YS, GV and NN; Writing: GB, TT and SH; Funding Acquisition: TT and SH; Resources: RG, AP and LK; Project Administration: SH; Supervision: TT and SH.

## Supplementary Figures

**Figure S1.**
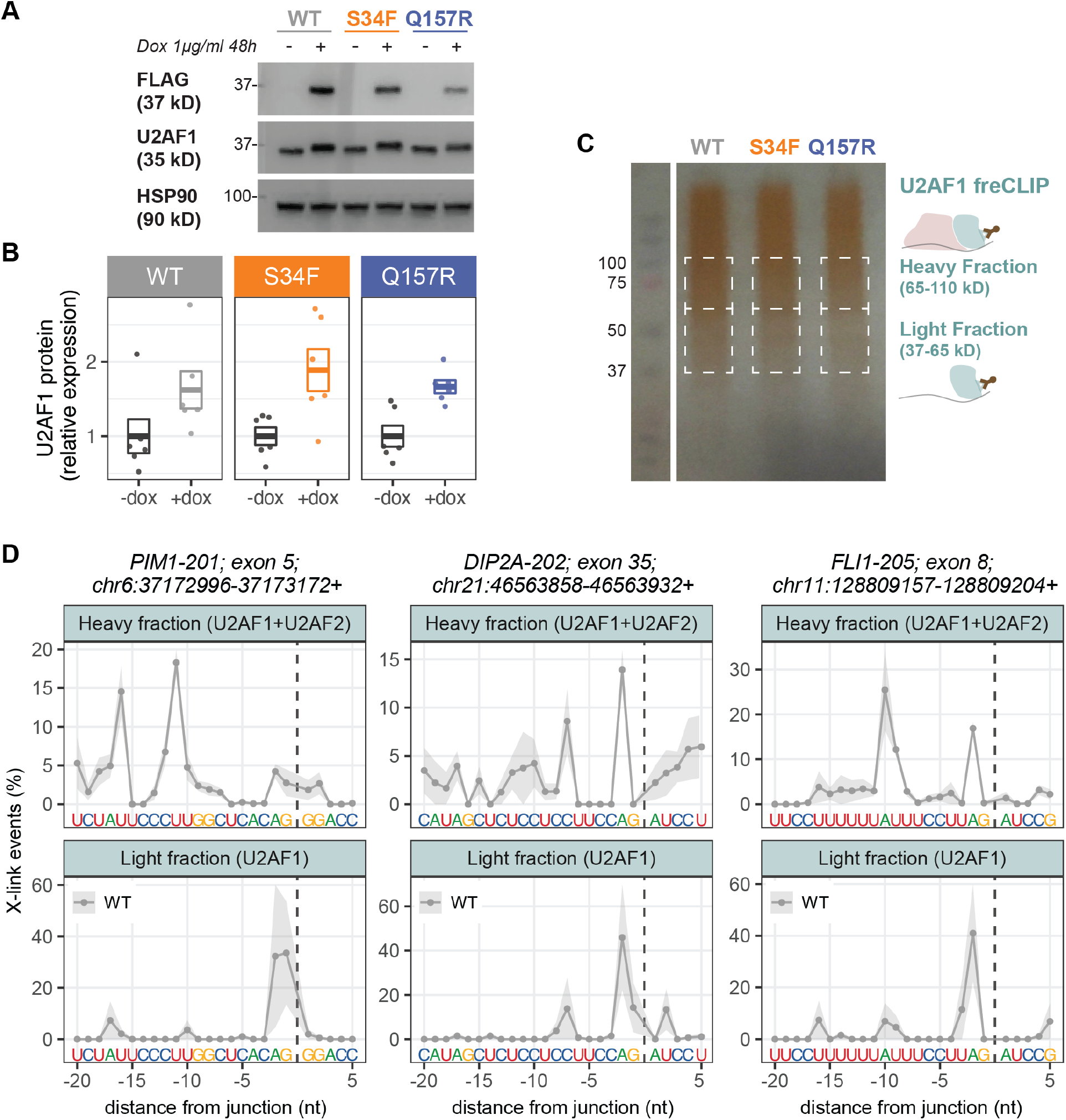
related to Figure 1. **(A, B)** Representative western blot **(A)** and quantification **(B)** of U2AF1 protein levels (WT, S34F or Q157R) before and after doxycycline induction. HSP90 was used as loading control. Mean±SEM among n=6 independent experiments are shown. **(C)** Representative U2AF1 freCLIP-seq size-selection to obtain light and heavy fractions. **(D)** Binding profile (mean±SEM) for light (n=3) and heavy (n=4) fractions of three representative single intron-exon junctions in U2AF1 WT freCLIP-seq. The 3’SS sequence of each junction is shown.

**Figure S2.**
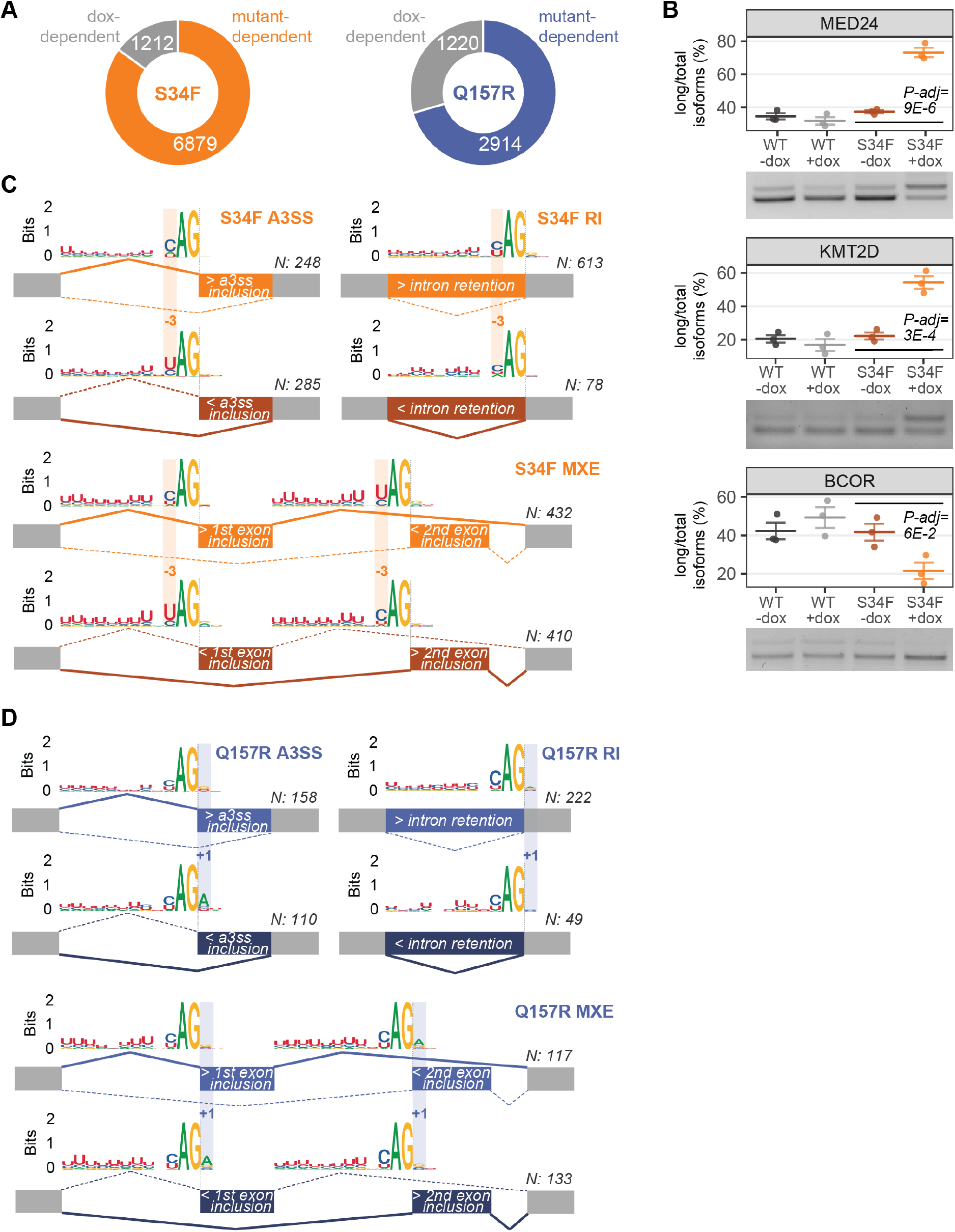
related to Figure 2. **(A)** Number of dox-dependent alternatively spliced events (grey section), comparing induced *vs* uninduced U2AF1 WT, that were removed to obtain the final list of differentially spliced events in S34F *vs* WT and Q157R *vs* WT. **(B)** Validation of alternative splicing level in three previously reported genes consistently identified by RNA-seq in U2AF1 S34F *vs* WT HEL cells. Representative gel images and corresponding quantification in n=3 biological replicates are displayed. P-adj: Tukey’s HSD test. **(C, D)** 3’SS sequence logos for alternative 3’ splice site events (A3SS), retained introns (RI) and mutually exclusive exons (MXE) in U2AF1 S34F *vs* WT **(C)** and in U2AF1 Q157R *vs* WT **(D)** HEL cells. Boxes highlight sequence-specific positions. N, number of alternative splicing events.

**Figure S3.**
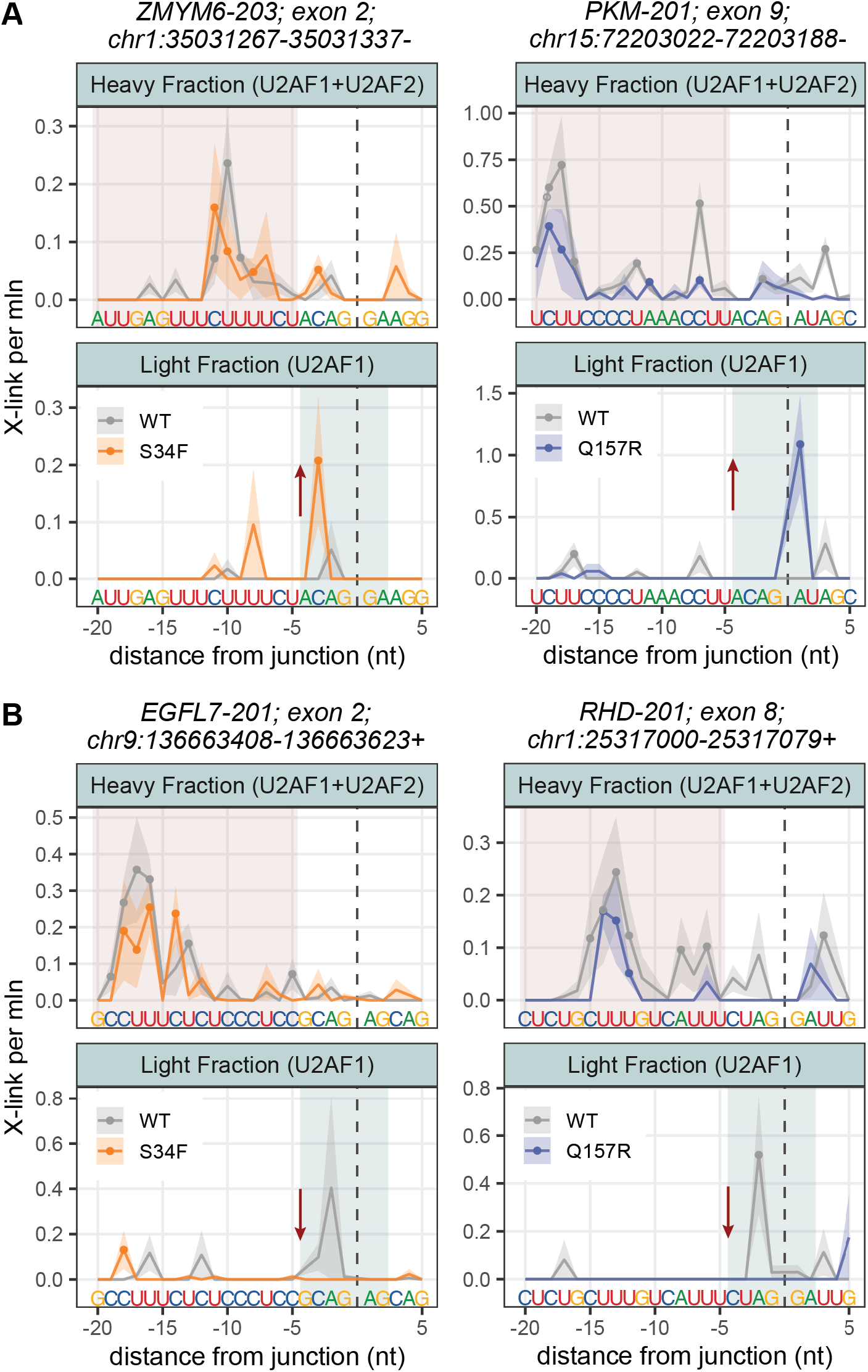
related to Figure 3. **(A, B)** Increased **(A)** *vs* decreased **(B)** mutant U2AF1 binding in four representative junctions. Binding profile (mean±SEM of the number of crosslinking events per million reads) for light and heavy freCLIP-seq fractions and 3’SS sequence are shown. Nucleotide positions with signal > 0 in all replicates are indicated by a dot (light and heavy fractions: WT, n=3 and n=4; S34F n=3 and n=3; Q157R, n=2 and n=2). Differential binding is defined comparing S34F or Q157R *vs* WT signals in −2;+4 light fraction region (U2AF1 monomer binding, light blue box) *vs* binding signals in −20;-5 heavy fraction region (pink box).

**Figure S4.**
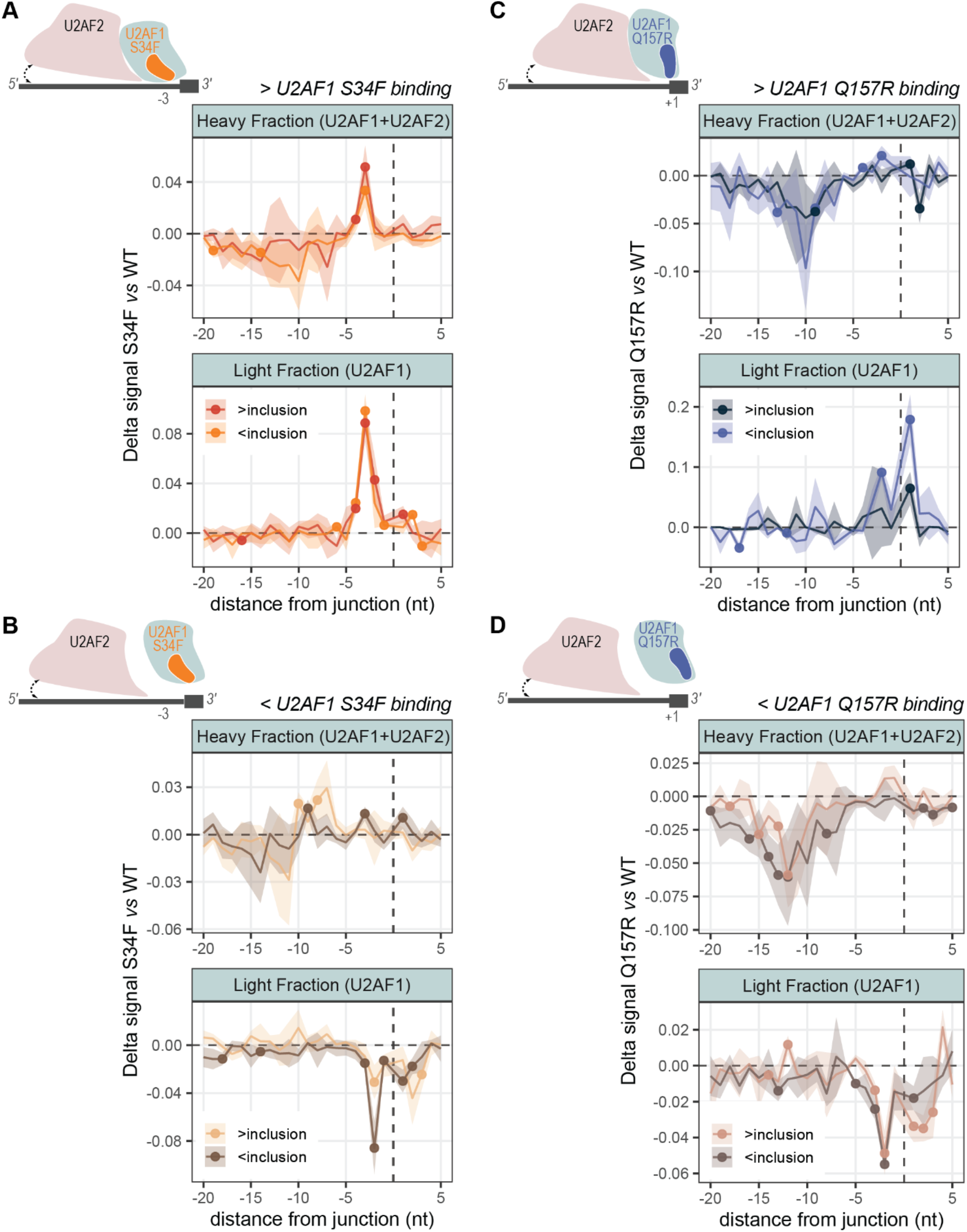
related to Figure 3. **(A-D)** Delta binding analysis between mutant and WT U2AF1 crosslinked sites per million in light and heavy freCLIP-seq fractions. Mean normalized for the size of each binding-splicing class ±SEM in each 3’SS position are shown. Positions characterized by significant binding change are indicated by a dot (P-value<0.05, one-tailed t-test). Junctions are grouped according to increased U2AF1 S34F binding **(A)**, decreased U2AF1 S34F binding **(B)**, increased U2AF1 Q157R binding **(C)**, and decreased U2AF1 Q157R binding **(D)**, with the corresponding schematic model representing the aberrant mutant binding in position −3 for S34F mutant or +1 for Q157R mutant and its influence on U2AF2 binding.

**Figure S5.**
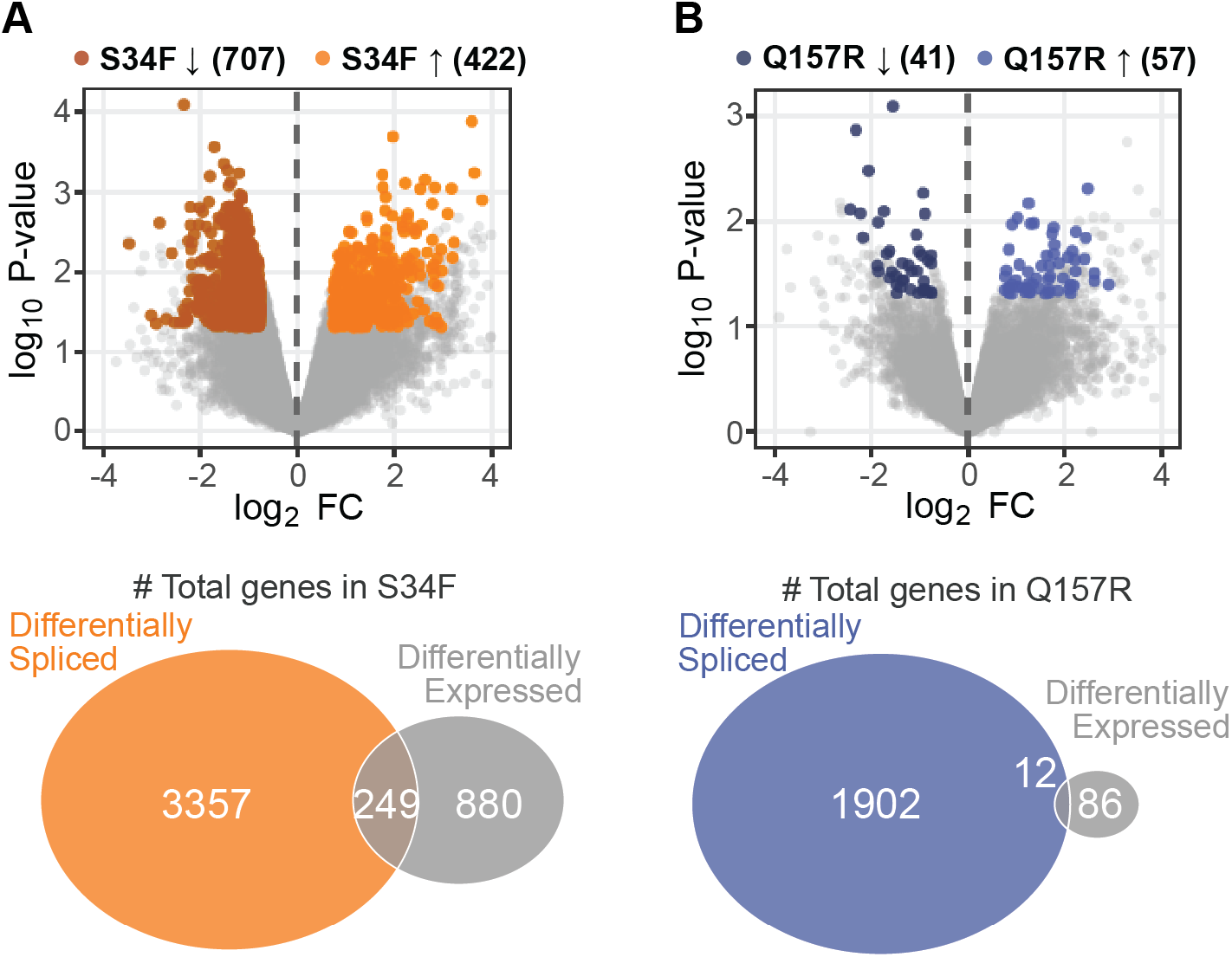
related to Figure 4. **(A, B)** Volcano plots showing fold change and P-value of differentially expressed genes in S34F *vs* WT **(A)** or Q157R *vs* WT **(B)** and comparison with differentially spliced genes. Colored and grey dots in volcano plots distinguish between significant and not significant differentially expressed genes (upregulated: log2 FC>0.75; P-value<0.05; downregulated, darker colors: log2 FC<−0.75; P-value<0.05; n=2 biological replicates per U2AF1 condition).

**Figure S6.**
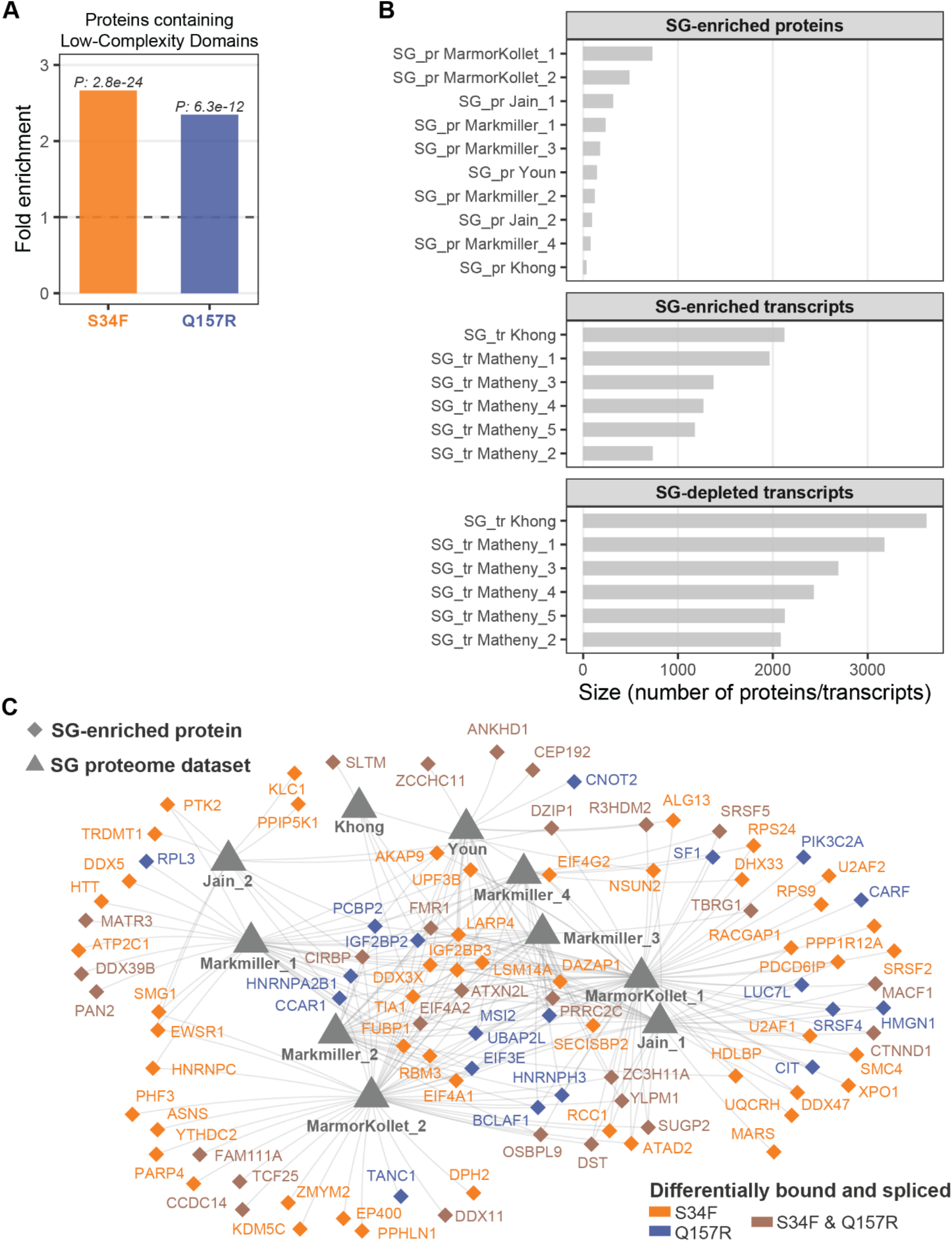
related to Figure 5. **(A)** Enrichment analysis of genes coding for proteins containing low-complexity domains considering differentially bound-spliced genes in U2AF1 S34F *vs* WT and Q157R *vs* WT HEL cells (Fisher’s exact test). **(B)** Number of stress granule-associated proteins or transcripts, enriched or depleted, in the meta-analysis of 16 published datasets (**Table S10**). **(C)** Network visualization of differentially bound-spliced transcripts whose corresponding protein is enriched in stress granules, according to multiple SG experimental datasets.

**Figure S7.**
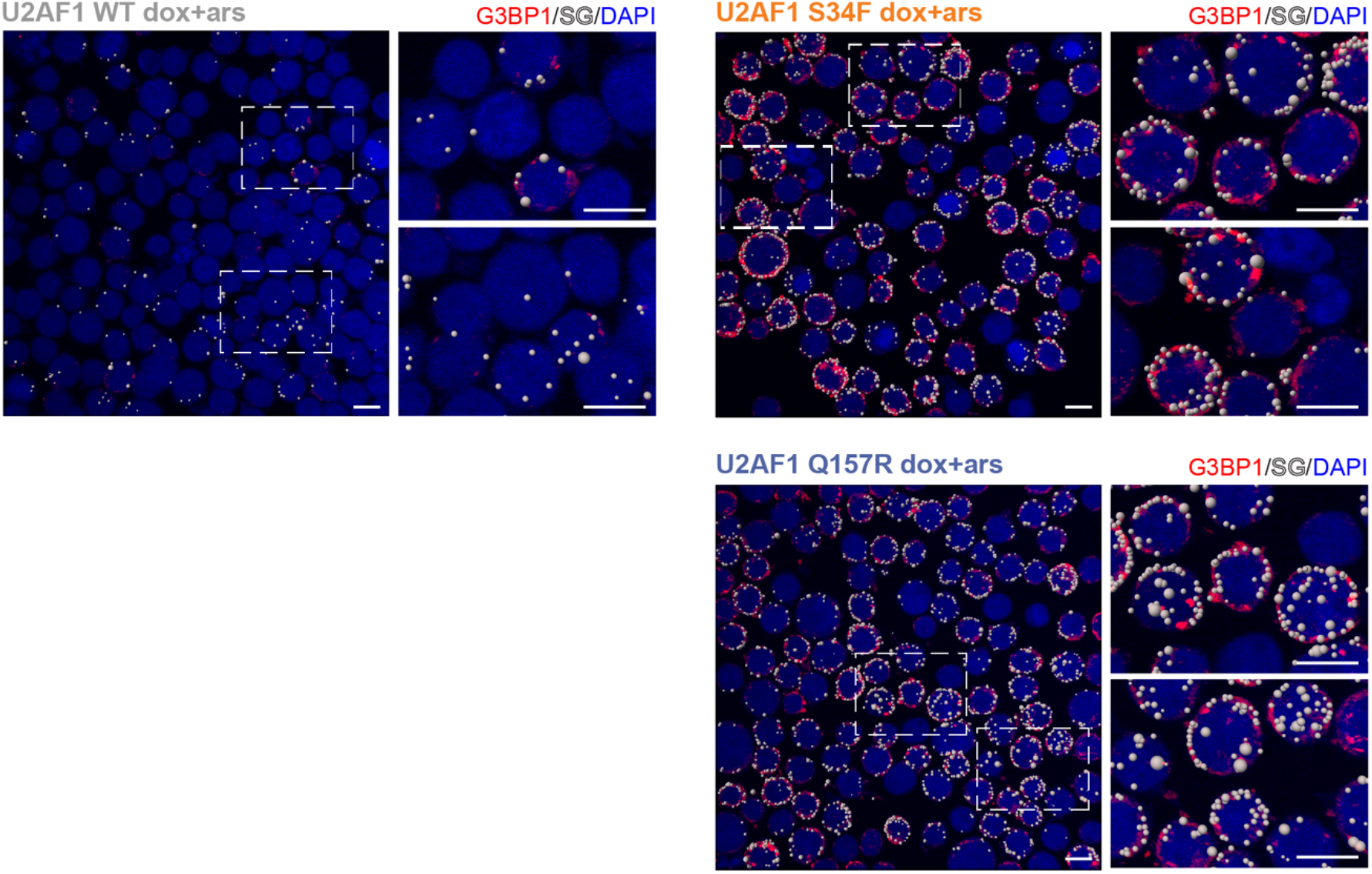
related to Figure 6. Representative IF images (scale bars, 10μm) of mutant and WT U2AF1 HEL cells. SGs identified by IMARIS (see Methods) are indicated by spheres.

**Figure S8.**
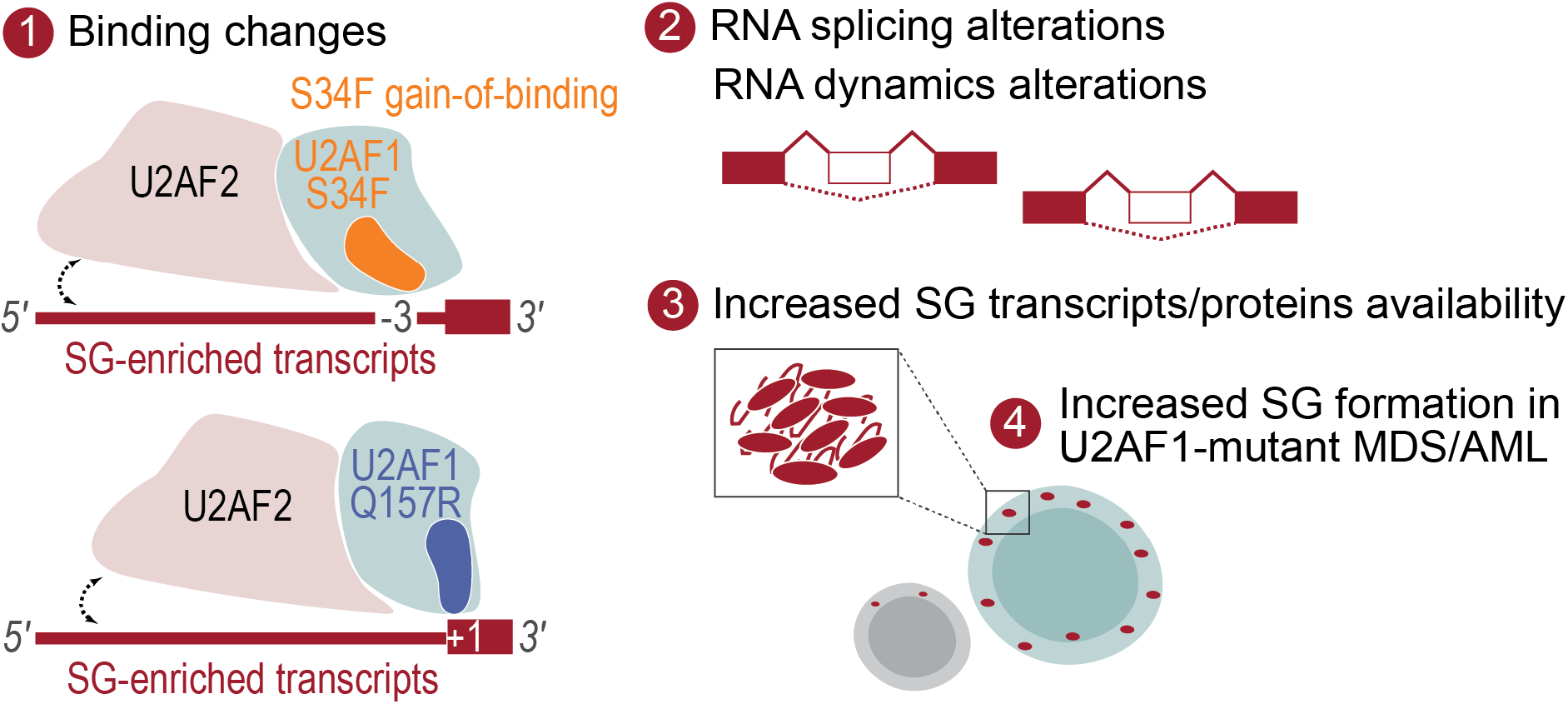
Model for stress granule perturbations in U2AF1-mutant myeloid malignancies.

## Supplementary Tables

***Table S1 related to Figure 1.*** eCLIP-seq metrics.

***Table S2 related to Figure 2.*** Alternative splicing events comparing S34F *vs* WT and Q157R *vs* WT U2AF1 HEL cells, after exclusion of dox-dependent events.

***Table S3 related to Figure 3.*** Differential binding events comparing S34F *vs* WT and Q157R *vs*

WT U2AF1 HEL cells.

***Table S4 related to Figure 3.*** Combined analysis of differentially bound and spliced genes comparing S34F *vs* WT and Q157R *vs* WT U2AF1 HEL cells.

***Table S5 related to Figure 4.*** Published datasets on U2AF1 mutation-dependent alternative splicing.

***Table S6 related to Figure 4.*** Comparative analysis of differentially spliced genes in U2AF1 S34F/Y and Q157R/P conditions in HEL RNA-seq data and in published datasets.

***Table S7 related to Figure 4.*** Functional annotation enrichment analysis of differentially spliced genes.

***Table S8 related to Figure 4.*** Functional annotation enrichment analysis of differentially bound genes.

***Table S9 related to Figure 5.*** Functional annotation enrichment analysis of differentially bound and spliced genes.

***Table S10 related to Figure 5.*** Published datasets on stress granule proteome and transcriptome.

***Table S11 related to Figure 6.*** IF imaging metrics for U2AF1 WT, S34F and Q157R HEL samples.

***Table S12 related to Figure 7.*** Patient characteristics.

***Table S13 related to Figure 7.*** IF imaging metrics for U2AF1 WT and S34F AML primary samples.

## Supplementary Files

***File S1-S2 related to Figure 6.*** Z-stack slice movie for representative U2AF1 S34F dox+ars (**S1**) and U2AF1 Q157R dox+ars (**S2**) images reported in Figure 6A.

